# Computational analysis of therapeutic potential for simplified Piper. spp- derived medicinal mixtures in anxiety, sleep, pain and seizure

**DOI:** 10.1101/2025.08.29.673131

**Authors:** B.G. Rice, B. Wooton, C. Jansen, J. Howard, J. Nixon, L. Ellis, A. Rogers, C.N. Adra, A.J. Stokes, S.A. Aporosa, A.L. Small-Howard, H Turner

## Abstract

Phytomedicines have played a vital role in traditional medical systems globally, particularly in providing culturally relevant and accessible healthcare solutions. *Piper methysticum*, known as Kava, is a traditional Pacific Island phytomedicine with clinically validated anxiolytic properties, primarily attributed to its Kavalactones. However, the biogeographically restricted distribution of *Piper methysticum* and the ecological and cultural concerns surrounding its widespread adoption highlight the need to explore alternative sources within the Piper genus. This study investigates whether other species within the Piper genus, used phytomedically in non-Pacific contexts, exhibit similar therapeutic efficacy for anxiety, stress, and related disorders including Post-Traumatic Stress Disorder (PTSD). We employed a computational approach utilizing a novel data platform of non-Western phytomedical pharmacopeias to analyze the secondary metabolomes of various *Piper* species. Network analysis and multidimensional data projections were used to compare the chemical composition and therapeutic indications of these species with those of *Piper methysticum*. Our findings suggest that while Kavalactones are predominantly unique to *Piper methysticum*, other *Piper* species also contain bioactive compounds associated with anxiolytic and stress-relieving effects. These results provide insight into the potential for culturally and biogeographically contextualized approaches to PTSD treatment, beyond the exclusive use of Kava, and lay the groundwork for future research into alternative phytomedicinal therapies within the Piper genus.

## Introduction

Kava (Piper methysticum G. Fost., Piper methysticum) is a culturally and pharmacologically significant plant in the Pacific region. *Piper methysticum* has increasing relevance for addressing complex neuropsychiatric and pain-related conditions that are underserved by current standards of care [1–4]. Kava is consumed by Pacific peoples in both ceremonial and less formal settings [5], and it has therapeutic indications in indigenous medical systems of Moananuiākea [6]. Its calming and restorative effects are aligned with therapeutically important indications including anxiety, post-traumatic stress disorder (PTSD), insomnia, pain, and also movement disorders associated with neurodegenerative diseases [7]. These symptoms frequently co-occur and exacerbate each other, forming a clinical constellation that is especially difficult to treat. For example, anxiety is a driver of insomnia [8], and sleep deprivation worsens both pain perception and emotional reactivity [9]. PTSD is characterized by a chronic hyperarousal state and often includes both pain and disrupted sleep [10,11]. Chronic pain is often co morbid with anxiety and PTSD [12]. Both pain and the dyskinesias and rigidity associated with Parkinson’s disease can lead to anxiety, impaired rest, and treatment complications [13,14]. The potential of Kava to act across these domains offers an unusual and potentially impactful therapeutic profile.

Standard pharmaceutical interventions remain inadequate across many of these indications. Benzodiazepines are effective anxiolytics with risks of tolerance, dependence, withdrawal, and cognitive impairment (and age-related decline in efficacy for elderly patients) [15–17].

Alternatives such as tricyclic antidepressants or gabapentinoids often have marginal efficacy and problematic side effects. Sleep aids (e.g., zolpidem, melatonin analogs) often fail in cases of chronic or trauma-related insomnia [18]. Pain management, especially for chronic, neuropathic, or inflammatory pain, is limited by the addictive nature of opioids [19]. In these contexts, Kava offers an attractive alternative or adjunctive strategy. Its clinical utility is supported by randomized controlled trials showing efficacy in anxiety, and empirical and traditional knowledge indicate effectiveness in sleep and pain. Importantly, Kava appears to induce a state of relaxed wakefulness rather than sedation [20,21], distinguishing it from conventional GABAergic medications and offering modulation of symptoms without impairment [22]. Its impact on sleep is to promote restorative sleep, which is critically important for PTSD and chronic pain. Adding to kava’s attraction is its high safety profile. Even when consumed in the large volumes typical of traditional use, research (e.g., WHO [23]) shows that kava is safe [24]. It is unlikely to cause balance issues or increase the risk of falls [22], and does not typically lead to liver toxicity or impair liver function [25]. This level of safety is reflected in how kava is regulated internationally, including regulation as ‘food’ in the Australia New Zealand Food Standards Code [26] and recent designation as Generally Regarded As Safe (GRAS) in the state of Hawai’i [27].

The entry of Kava into mainstream clinical use faces logistical, ecological, and epistemological challenges. First, there are supply chain issues: Biogeographically, *Piper methysticum* is a cultural keystone species unique to the Pacific where island nations face increasing vulnerability to climate change and economic instability. Kava production is often reliant on smallholder agricultural systems with limited scalability [28]. Past surges in international demand have triggered boom-bust [29,30] cycles, leading to ecological damage, land dispossession, and concerns over biopiracy and unequal benefit-sharing [31]. This challenge creates the need to explore alternative supply chain options that will not have negative implications for the Pacific owners of kava. Second, there are formulation issues: From a regulatory and quality control standpoint, commercial products containing *Piper methysticum* are variable, ranging from aqueous traditional preparations to solvent-extracted capsules. The latter often fail to replicate traditional formulations in either composition or effectiveness, and the type of simplification to one or two constituents that is a pre-requisite for regulatory approval by agencies such as FDA will likely also face failure to recapitulate full therapeutic effects [32,33]. Outside culturally-informed practice [20,34], Kava remains a nutraceutical in countries like the US, resulting in essentially unregulated preparation and associated risks to patients. Abstraction from traditional preparation has also resulted in pharmacovigilance alerts such as hepatotoxicity which is not associated with traditional usage, but has previously been observed in culturally-decontextualized consumption [20]. These challenges create the need to rationally design Kava-based or -inspired formulations that are reduced in complexity (i.e. Minimal Essential Effective Formulations, MEEF) that recapitulate effectiveness and can viably navigate drug discovery and approval pipelines. Third, there are mechanistic questions: The primary pharmacological activity of Kava is attributed to kavalactones, which are low-affinity positive allosteric modulators of GABA-A receptors. However, the complexity of the Kava secondary metabolome and the breath of its impacts on physiology leave open the question of whether kavalactones are necessary, sufficient, or merely contributory agents to all its therapeutic effects. This issue creates a need to reliably match secondary metabolites (including but not limited to kavalactones) to specific therapeutic benefits.

This paper uses computational methods to begin addressing these challenges. It explores the identification of *Piper methysticum* alternatives within the broader *Piper* genus based on traditional usage in target disorders of interest (anxiety, pain, sleep, movement/seizures), linking those relationships to chemical composition with a focus on establishing the primacy, or otherwise, of the kavalactones. A proof-of-concept is provided using a novel phytomedical data platform (PhAROS™) that curates ethnomedical, phytochemical, and pharmacological data across Global Integrative Medical Systems (GIMS). Using chemical composition data from multiple studies, we constructed an integrated metabolome of *P. methysticum* and assessed its extent of overlap with other *Piper* species was assessed. This revealed that while kavalactones are relatively unique to Kava, other *Piper* species are rich in alkaloids, phenylpropanoids, and terpenes with strong associations to the same therapeutic indications for anxiety, pain, sleep, and movement/seizures. A zebrafish behavioral model was then used to validate the anxiolytic potential of select phytochemicals, both from Kava and from other *Piper* spp. Notably, yangonin, a non-sedating component of *P. methysticum*, and piperine, a bioactive from *P. nigrum* demonstrated stress-response modulation without inducing motor suppression. This favorable profile is not achieved by all kavalactones. Druggability analytics in the PhAROS^TM^, including Lipinski’s Rule of Five, natural product-likeness scores, and weighted QED indices, were used to rank compounds suitable for future phytopharmaceutical development. These analyses revealed that several non-kavalactone candidates not only mimic therapeutic functions but are also tractable for formulation into standardized, regulatory-compliant products. The combination of trans-cultural, trans-species, and strategic dereplication into one computational workflow, combined with a pathway to validated in vivo prescreening, offers a route to MEEF that captures but simplifies the inherent polypharmacy of phytomedicines such as Kava.

Finally, three strategic pathways are identified for the development of Kava-derived and Kava-inspired medicines. The first maintains maximal fidelity to traditional preparation and use, preserving Kava’s full chemical and relational architecture (e.g., naturalist Kava use such as that advanced in the *Kava-talanoa* framework for PTSD). While this pathway honors cultural sovereignty and likely delivers the most authentic therapeutic effect, it presents challenges for adoption, standardization, reproducibility, and regulatory approval. At the opposite end lies a reductionist strategy involving the synthesis or purification of single kavalactones or analogs. This may facilitate pharmacological clarity and scalability but risks the loss of efficacy that is founded in polypharmacy, and its cultural disconnection is potentially extractive [35,36]. The third ‘middle-road’ approach explored here aims to define a minimal essential polypharmaceutical profile that retains synergy and integrative effects, that maintain cultural fidelity but are more viable for regulatory development. This middle road is rendered feasible by PhAROS^TM^ as a novel computational space that is biogeographically and culturally agnostic, and de-siloed from these conventional constraining boundaries.

## Materials and Methods

### Construction of the PhAROS Data Platform

#### Data Acquisition

Global Integrative Medical Systems (GIMS) datasets were pulled into a data warehouse via site download options or web data extraction from Encyclopedia of Traditional Chinese Medicine (ETCM), Ethnomedicinal Plant Database (ETM-DB), Indian Medicinal Plants, Phytochemistry And Therapeutics (IMPPAT), South African National Chemical Database (SANCDB), Kampo Medicine Database (KampoDB), Kyoto Encyclopedia of Genes and Genomes (KEGG), Northern African Natural Products Database (NANPDB), Taiwanese Indigenous Plant Database (TIPdb), Northeast Asian Traditional Medicine (TM-MC), Traditional Chinese Medicine Integrative Database (TCMID), Bioactive Phytochemicals Molecular Database (BioPhytMol), Medicinal Plant Server (MedPServer), Korean Traditional Knowledge Portal (KTKP), and Pharmacological Database of Korean Medicinal Plants (PharmDB-K). Each traditional medicine system dataset was preprocessed to include most of the following features: System of origin (TXM), ingredient combination (Formula), ingredient organism scientific name (Species), compounds associated with each species (Compound), compound identifier (CID), and usage (Indications)

#### Data Enrichment

The GIMS datasets were enriched with plant-phytochemical lists from Dr. Duke’s Phytochemical Database [37,38] and the compounds were further enriched with Pubchem information to include CID, INCHIKEY, and Classification. Missing compound classifications were filled in using Classyfire (Edmonton, Alberta, Canada) [39]. Country codes were matched to appropriate ISO 3166 Codes for mapping. Protein targets were matched to compounds via CHEMBL identifiers and bioassays.

### Indication dictionary construction

The indication dictionary delineates the search terms and truncations used to interrogate PhAROS with the aim of identifying data fields linked to certain disorders.

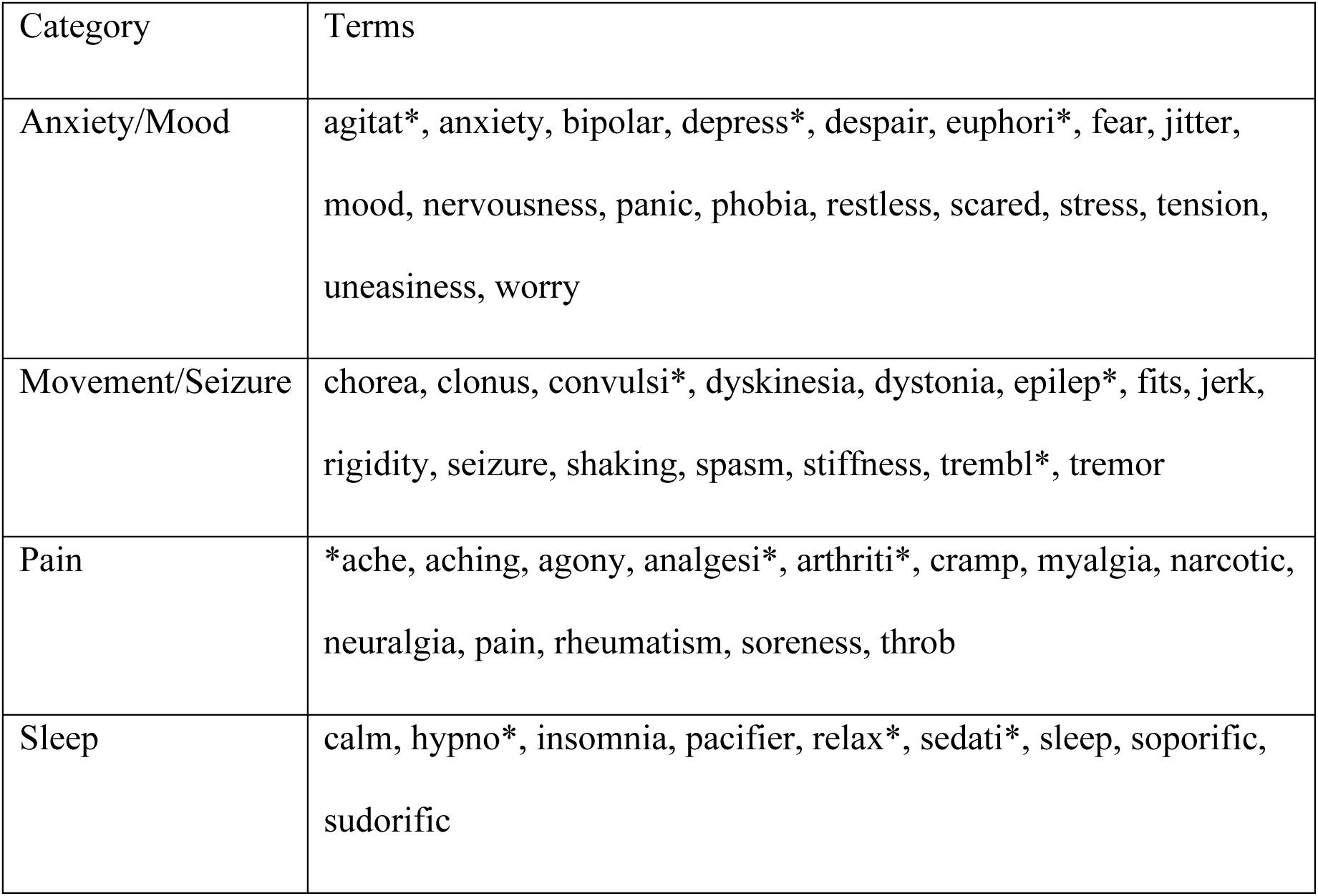

| Category | Terms |
| --- | --- |
| Anxiety/Mood | agitat*, anxiety, bipolar, depress*, despair, euphori*, fear, jitter, mood, nervousness, panic, phobia, restless, scared, stress, tension, uneasiness, worry |
| Movement/Seizure | chorea, clonus, convulsi*, dyskinesia, dystonia, epilep*, fits, jerk, rigidity, seizure, shaking, spasm, stiffness, trembl*, tremor |
| Pain | *ache, aching, agony, analgesi*, arthriti*, cramp, myalgia, narcotic, neuralgia, pain, rheumatism, soreness, throb |
| Sleep | calm, hypno*, insomnia, pacifier, relax*, sedati*, sleep, soporific, sudorific |

### GBIF Plots

The Global Biodiversity Information Facility (GBIF) database was used to map the global distribution of *Piper* species. Occurrence data at the species level was downloaded from the database [40]. Entries flagged for duplication or quality issues were excluded to ensure data integrity. A map of species occurrences was generated and downloaded directly from the GBIF interface. The final map was rendered in R and annotated with species names and counts per region, providing a global overview of documented *Piper* diversity as recorded in publicly accessible biodiversity repositories.

### Data Analysis

Exploratory data analysis was conducted via the following python packages: pandas, numpy, matplotlib, and seaborn. Plotly, biophyton, networkx and chembl_web_client were additional Python packages used in the frequency analysis. Circos [41] was used to construct chord diagrams. The python package wordcloud was used to construct word clouds, utilizing built-in tools such as STOPWORDS to filter out common words and ImageColorGenerator to apply color schemes based on input images.

### Druggability analyses

Standard InChIKeys (fixed-length, SHA-256 hashed counterpart of the standard International Chemical Identifier) and canonical SMILES (Simplified Molecular Line Entry System) strings for all compounds in the PhAROS data platform were obtained from the PubChem PUG-REST API [42,43]. These InChIKeys were then used to join compounds in PhAROS™ with the set of compound properties provided by the ChEMBL v32 database, to obtain NP-likeness scores, Lipinski’s rule of 5 violations, and weighted QED scores [44]. For those compounds not present in ChEMBL, the open source RDKit cheminformatics software was used to compute the number of Lipinski’s rule of 5 violations and weighted QED scores [45]. To impute missing NP-likeness scores for compounds without a ChEMBL cross-reference, a statistical imputation technique leveraging the scikit-learn package’s built-in RandomForestRegressor model [46] was used. The hyperparameter settings of the model were: max tree depth of 80, and number of trees in the ensemble of 150. The features used as predictors to train the imputation model were as follows: 2048-bit length ECFP4 2d fingerprint features, molecular weight, number of hydrogen bond acceptors, number of hydrogen bond donors, polarizable surface area, number of rotatable bonds, and ALogP. The model was trained and evaluated on the set of 14,961 PhAROS compounds with known NP-likeness scores obtained from ChEMBL v32, at an 85%/15% train/validation split. The imputation model achieved an R^2^ score of 0.9412 and an MSE (mean squared error) loss of 0.0772 on the validation set. The performance of the imputation model is summarized in Supplemental Figure A. This trained model was then used to impute the NP-likeness score for the remaining 34,540 compounds in the PhAROS™ database without a ChEMBL cross-reference. As a result, 69.76% of the compounds in PhAROS™ have imputed NP-likeness scores.

Having obtained weighted QED, NP-likeness, and Lipinski’s rule of 5 scores for all compounds, we then visualized the distributions of these metrics for four non-disjoint sets of compounds: (1) all natural compounds contained in the PhAROS™ data platform, (2) all natural compounds reported in the *Piper methysticum* secondary metabolome, (3) all natural compounds found in other *Piper* spp. and associated with indications of interest, and (4) the eight natural compounds tested in our in vivo pilot study. The distributions of the metrics for these four sets of compounds are presented in Figure 11.

### Fish husbandry

Adult zebrafish (*Danio rerio*) were maintained according to standard animal care protocols [47] in accordance with the Canadian Council of Animal Care (CCAC) guidelines. Adult AB/Tubingen zebrafish were housed on a re-circulating aquatic system at 28.5 ± 1 °C, pH 7.0–7.2 on a 14:10-h light:dark schedule. Age matched fertilized embryos were housed (Pentair Aquatic Eco-system, Apopka, FL, USA) in nursery baskets (< 200 embryos per basket) in 3-L tanks (Tecniplast, Buguggiate, VA, Italy), with matching conditions to adults. Larvae were removed from the re-circulating system using HEPES buffered E3 (HE3) medium (5 mM NaCl, 0.17 mM KCl, 0.33 mM CaCl2-2H2O, 0.33 mM MgSO4-7H2O, 10 mM HEPES, pH 7.2) at 120 h post-fertilization (hpf) to be used in experiments that day.

### Assessing anxiolytic potential from acute exposure to individual phytochemicals using a model of anxiety-like larval behavior [48]

At 120 hours post fertilization (hpf), zebrafish larvae were transferred to a 48 well plate (Thermo Scientific) with 1 larva per well in 450 μL of HEPES buffered E3 (HE3) medium (5 mM NaCl, 0.17 mM KCl, 0.33 mM CaCl2-2H2O, 0.33 mM MgSO4-7H2O, 10 mM HEPES, pH 7.2) using a micropipette and acclimated for 2 hours at 28 °C in a light incubator. Pure compounds were stored as 0.5 mg/mL (Yangonin only) and 1 mg/mL in MeOH stock solution at 4 to - 20°C. Working solutions were prepared fresh each day at a 10x concentration. 50 μl of the 10x solution was pipetted into each well to reach the final concentration. Each experimental replicate was comprised of 12 larvae per extract concentration. Following the addition of the phytochemical solution each plate was placed directly into the automated behavioral tracking system and exposed to our standardized behavioral assay. The 2-hour standard behavioral assay consisted of 1.5 hours of consistent light followed by alternating 5-minute periods of light or dark for 30 minutes. Following the behavioral assessment, larvae were visually scored for any abnormalities. Dead, necrotic or visually affected larvae were removed from the behavioral analysis. The patterns of acute behavior were assessed using the Noldus EthoVision XT16-17 software. The standard measure of activity was the total distance traveled quantified into 60-second bins. The assay was completed a minimum of 2-3 times for each concentration of phytochemical (n=24-36) and activity levels measured were compared to controls using 2-WAY ANOVA followed by a Dunnett’s multiple comparison test where p<0.05. Impacts on larval behavior were assessed by comparing the average distance traveled during the first 90 minutes and the dark response (calculated by subtracting the preceding 5 minutes of movement in the light from the first 5 minutes in the dark) to controls using one-way ANOVA followed by a Dunnett’s multiple comparison test, with significance set at p<0.05.

### Assessing anxiolytic potential from exposure to individual and mixtures of phytochemicals using a thigmotaxis model of larval behavior

Another model of anxiety/stress-like behavior utilized in this project is adapted from previous studies of zebrafish larval anxiety/stress using thigmotaxis (wall hugging) as the main indicator of stress [48–51]. At 48 hours post fertilization (hpf), zebrafish embryos were transferred to a 24 well plate with 900 μL of buffered media using a pipettor, sealed with film and placed in a light cycling incubator (28°C) until 120 hpf. Published controls used were caffeine and diazepam. Pure compounds for the current project were stored as 0.5 or 1 mg/mL in MeOH stock solution at 4 to -20°C. Phytochemical working solutions were prepared fresh each day at a 10x concentration. 100 μl of the 10x phytochemical or carrier control solution was pipetted into each well to reach the final concentration. Once dosed the plates were transferred directly into the automated behavioral tracking system and exposed for either 3, 6 or 10 minutes in the light and 4 mins in the dark. Dead, necrotic or visually affected larvae were removed from the behavioral analysis. The patterns of behaviors were assessed using the EthoVision XT16-17 software. Each experimental replicate was comprised of 12 larvae per extract concentration and the assay was completed a minimum of 2-3 times for each concentration of phytochemical compound or mixture (n=24-36). Behavior was only measured in the 4-minute dark period where fish would be exhibiting scototaxis (increased movement as a stress response) which would highlight any larvae that were not moving and could be eliminated from the analysis. Thigmotaxis was calculated as the % of time spent in the outer zone compared to the total time of the experiment and was compared to controls using ANOVA and multiple comparison’s test (p<0.05).

### Definition of Kava for the purposes of this study

For this study, *kava* is defined not merely as the beverage produced by mixing the ground roots of *Piper methysticum* with water, nor as commercial derivatives such as extracts, capsules, or beverages containing *Piper methysticum*. These commercial forms vary in composition and safety profile and may include additives such as kratom (*Mitragyna speciosa*) [52]. In alignment with Pacific traditional knowledge, *kava* is understood as a holistic cultural construct that extends beyond physical form to encompass ceremonial and relational contexts in which it is consumed [53], including the practices of *talanoa* (a dialogic and relational mode of engagement that integrates physical, spiritual, ancestral, and communal dimensions [54,55]. The concept of *talanoa-vā* further emphasizes the spatial and temporal relationships maintained through trust, mutual understanding, and ongoing dialogue [56–58]. Accordingly, throughout this research, the term *kava* is used in its culturally grounded sense, reflecting its embeddedness in Pacific epistemologies and relational systems. Use of the term to refer solely to the plant or its isolated chemical constituents is considered an oversimplification and misrepresentation of its traditional meaning.

## Results

### Comparative biogeographical and therapeutic distributions of *Piper spp*. and *Piper methysticum*

We are interested in exploring whether other species within the Piper genus, beyond *Piper methysticum* (Kava), offer viable therapeutic alternatives for the treatment of PTSD, anxiety, movement and sleep disorders, potentially providing strategies to mitigate the biogeographic, cultural, and supply chain limitations associated with a sole focus on Kava. We used the Global Biodiversity Information Facility (GBIF) database [40] spanning the years 1838 to 2019 to assess documented occurrences of *Piper spp.* and *Piper methysticum* worldwide. Digitized occurrences in this database include both growing specimens in situ and voucher specimens transported from their endemic location. Figure 1A illustrates the worldwide distribution of Piper species, with text boxes highlighting specific *Piper* species utilized in traditional medicine systems. Between 1600 and 2022, there were 377,334 occurrences of the various *Piper* species recorded in GBIF, 212,858 of which were georeferenced to 169 unique countries/areas. The number of species associated with traditional medicine in different regions is outlined: Americas (70 species), Africa (14 species), Europe (3 species), Oceania (15 species), and Asia (76 species). Figure 1B illustrates the distribution of 608 digitized occurrences of *Piper methysticum* G. Forst. recorded in GBIF. Of these occurrences, 121 were georeferenced with the majority concentrated in Oceania. Specific locations include French Polynesia (24 occurrences), Micronesia (19), Hawaii (17), Vanuatu (13) [6], Papua New Guinea (5), Fiji (5), Indonesia (4), Cook Islands (3), Niue (2), Cote d’Ivoire (1), Tonga (1), and Samoa (1). The occurrences observed along the west coast of the continental US are documented as preserved specimens held by the California Department of Food and Agriculture and do not support claims that *Piper methysticum* has cultural keystone species status on Continental USA. Occurrences in Africa are preserved specimens housed in Les spermaphytes de l’Herbier du Centre National de Floristique de Côte d’Ivoire. These data support the idea that *Piper methysticum* is largely endemic to Oceanic Pacific islands but other species in the family have a far wider global distribution.

**Figure 1.**
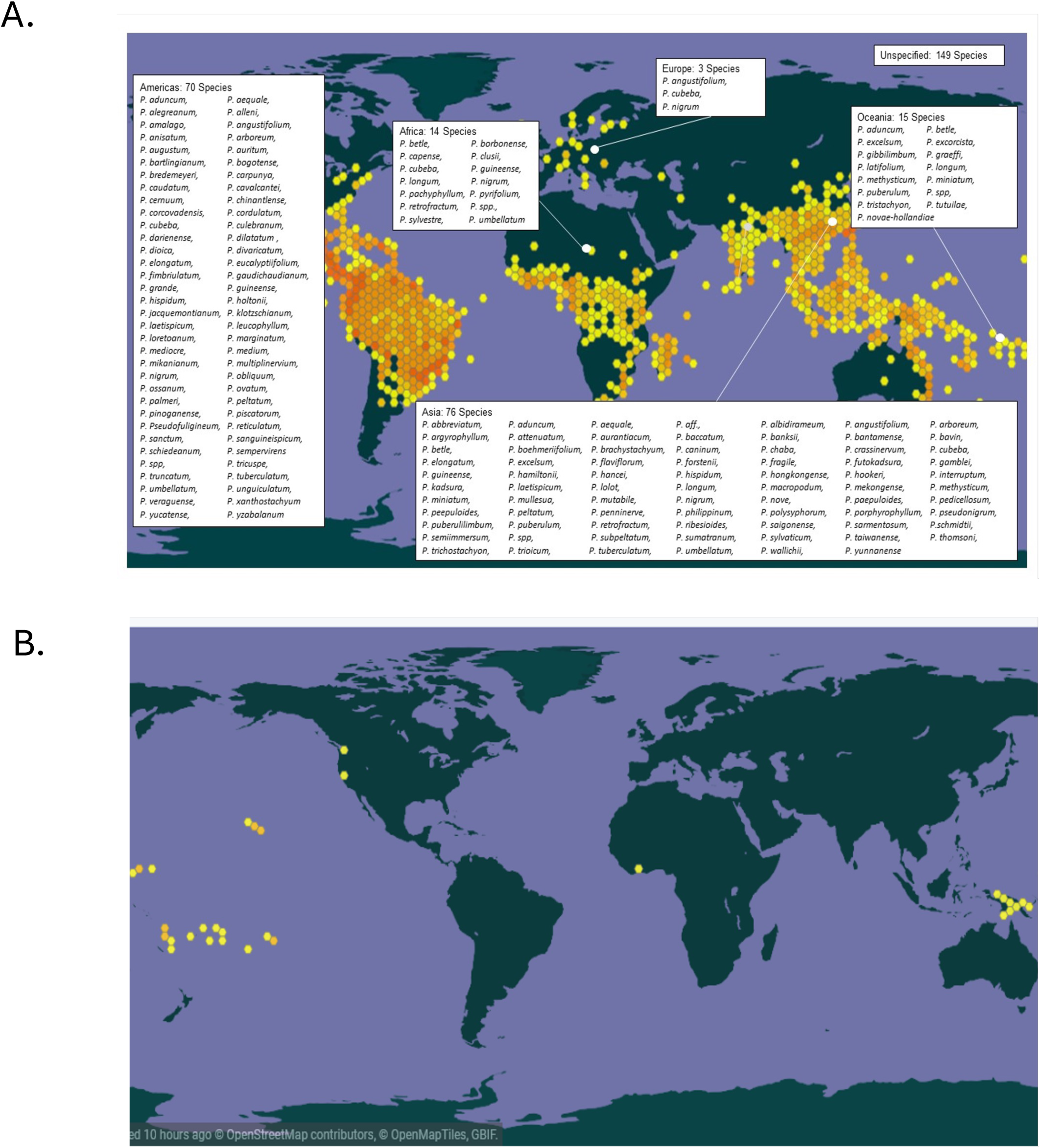
Global distribution of *Piper* species and geographic specificity of *Piper methysticum*. **A. Map showing the worldwide distribution of *Piper* spp. documented in the Global Biodiversity Information Facility (GBIF) from 1600 to 2024.** Hexagonal density overlays indicating frequency of occurrence. Text boxes highlight the number and identity of species used in traditional medicine within each region. A total of 377,334 occurrences were recorded, with 212,858 georeferenced across 169 countries. Regional representation of medicinally used *Piper* species includes: Americas (70 species), Asia (76 species), Oceania (15 species), Africa (14 species), and Europe (3 species). **B. Georeferenced occurrences (n = 121) of *Piper methysticum* G. Forst.** extracted from the same GBIF dataset. The species is largely endemic to Oceania, with confirmed occurrences concentrated in regions such as French Polynesia, Micronesia, Hawaii, Vanuatu, Papua New Guinea, and Fiji. Additional occurrences likely reflect preserved/voucher specimens rather than in situ populations (e.g., California Department of Food and Agriculture, *Les spermaphytes de l’Herbier du Centre National de Floristique de Côte d’Ivoire*).

We next assessed the medical as opposed to biogeographic distribution of the *Piper* species using a novel phytomedicine analytics tool. We constructed a Data Platform (termed PhAROS^TM^) and a data analytics workflow termed In Silico Convergence Analysis (ISCA) (see Methods and Table I) to provide for analysis of the relationships between plant species, indications and chemical constituents across a range of phytomedicine pharmacopeias in various geographical and cultural settings. Figure 2A shows a schematic of the data platform assembly and content.

**Figure 2.**
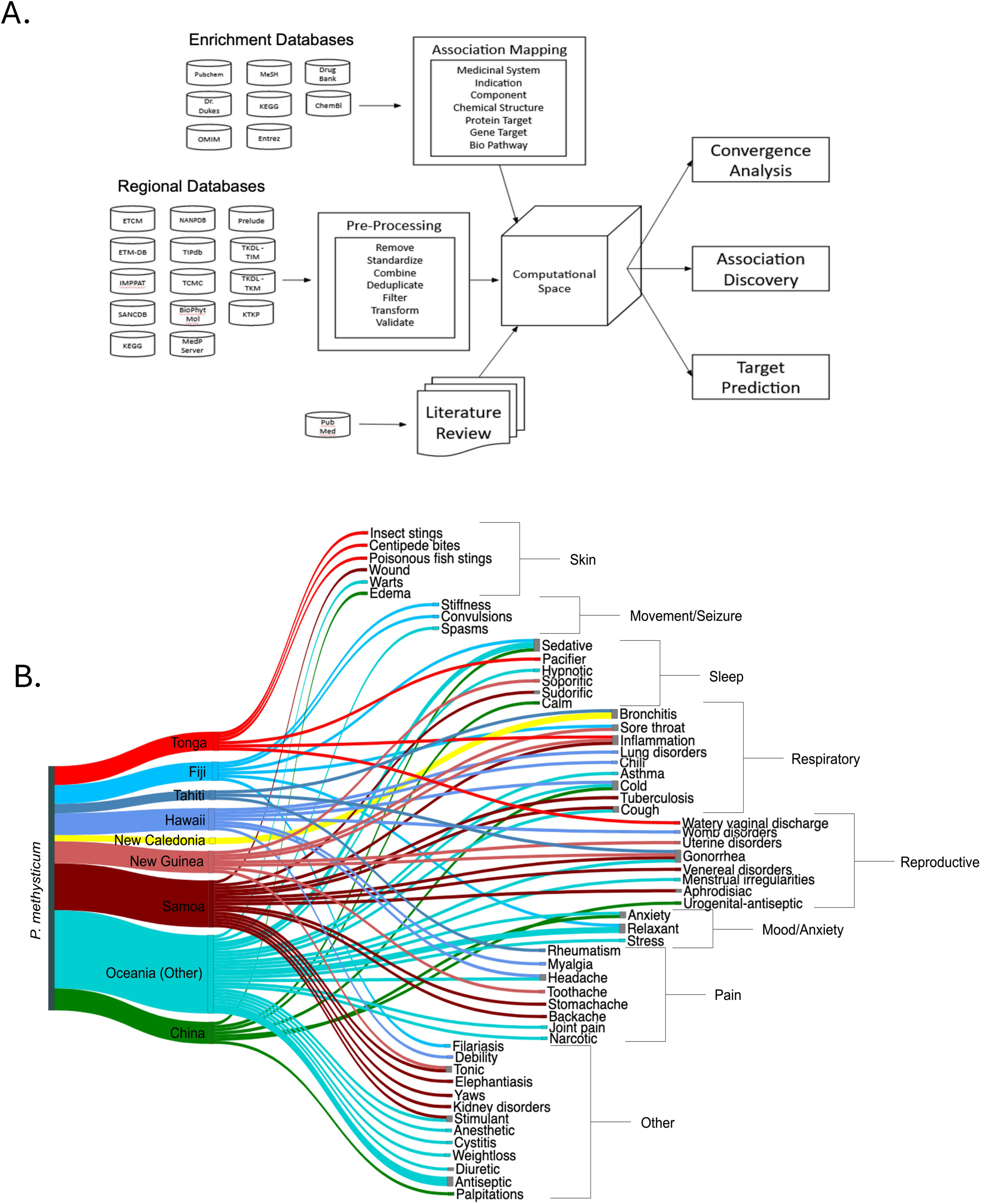
Computational framework and global indication mapping for *Piper methysticum* across traditional medicine systems. **A.** Schematic overview of the PhAROS Data Platform architecture and *in silico* convergence analysis (ISCA) workflow. The system integrates compound, indication, and ethnomedical data from diverse Global Integrative Medical Systems (GIMS), alongside chemical and pharmacological enrichment databases. Data undergo preprocessing for harmonization and deduplication before being integrated into a computational space enabling association mapping, convergence analysis, target prediction, and discovery of indication-chemical relationships. Outputs include chemical-indication networks, bioactivity predictions, and hypothesis generation for phytomedical efficacy. **B.** Sankey diagram visualizing the therapeutic indications of *Piper methysticum* (Kava) as documented in GIMS pharmacopeias. Indications span a broad range of physiological systems, including anxiety/mood, sleep, movement/seizure, respiratory, pain, reproductive, and skin-related conditions. Several regions employ *P. methysticum* for calming effects (e.g., sedative, pacifier, relaxant) and reproductive disorders, with respiratory ailments (e.g., bronchitis, asthma, cough) also frequently treated.

We constructed a Sankey diagram in Figure 2B to visualize the indications treated by *Piper methysticum* in GIMS formulations across geographical regions. *Piper methysticum* is predominantly used in traditional medicine in Pacific islands and Pacific diasporic communities in wider Oceania [32,59–61], with additional incorporation into medicinal practices in parts of Asia (China). Notably, each region uses the plant for calming effects. Reproductive disorders and respiratory-related diseases are common use cases.

As described above [1–4], *Piper methysticum* is expanding both in nutraceutical usage, as a Western allopathic pharmaceutical and in naturalistic usage outside the Pacific. Areas of emerging interest for Kava therapeutics include Anxiety (including Generalized Anxiety Disorder, GAD and Social Anxiety Disorder, SAD), PTSD, stress, insomnia and sleep disorders [20,62,63], as well as pain [64–66]. Within this grouping of potential therapeutic uses there is interest in possible impact on disorders such as Parkinson’s Disease. In PD patients, Kava may be a potential therapeutic for anxiety and sleep quality issues, and there is some interest in potential neuroprotective effects of kava component [67–70]. Kava is also traditionally used as a muscle relaxant which may provide potential relief in the dyskinesia associated with PD [71], although this has not been explicitly studied. We note also that many anti-seizure medications work by enhancing GABAergic activity, suggesting a potential for Kava to have some influence on seizure activity [72]. Based on these potential use cases for kava, we constructed term dictionaries (see Methods) that group four main indication categories to try to capture *Piper spp.* usage in GIMS that would encompass instances of medical usage in in Western and Pacific settings. These four selected categories are Mood/Anxiety, Sleep, Pain and Movement/Seizure. We then used these categories to interrogate PhAROS™ and assess whether *Piper spp*. other than *Piper methysticum* are used to address these indications.

### Representation of Mood/Anxiety, Sleep, Pain and Movement/Seizure disorder categories in indication dataset for use of Piper spp. across GIMS

We performed a multidimensional data analysis and visualization to assess indications for *Piper spp.* across multiple GIMS phytomedical pharmacopeias. We used a Circos plot [41] (Figure 3A) to visualize relationships between *Piper* species and the four major indication categories of interest. The Circos plot visualizes the frequency of usage of Piper species in various formulas across different countries. The visualization shows that *Piper spp.* other than *Piper methysticum* have significant representation in formulations for each of these major categories. *Piper nigrum, Piper longum*, and *Piper kadsura* are most frequently employed in formulations related to the categories of interest. *Piper borbonense, Piper cubeba, Piper guineense, Piper kadsura, Piper longum, Piper methysticum, Piper nigrum, and Piper umbellatum* are used in formulas addressing all four major indication categories and an additional nine species show associations with two or more indications. By percentage, *Piper methysticum*, *Piper kadsura*, and *Piper borbonense* are over-represented in formulations that treat these major disorder categories despite their relatively low frequency of representation in the whole Piper dataset. Interestingly, pain emerges as a predominant indication for *Piper spp*., dominated by *Piper nigrum* and *Piper longum*.

**Figure 3.**
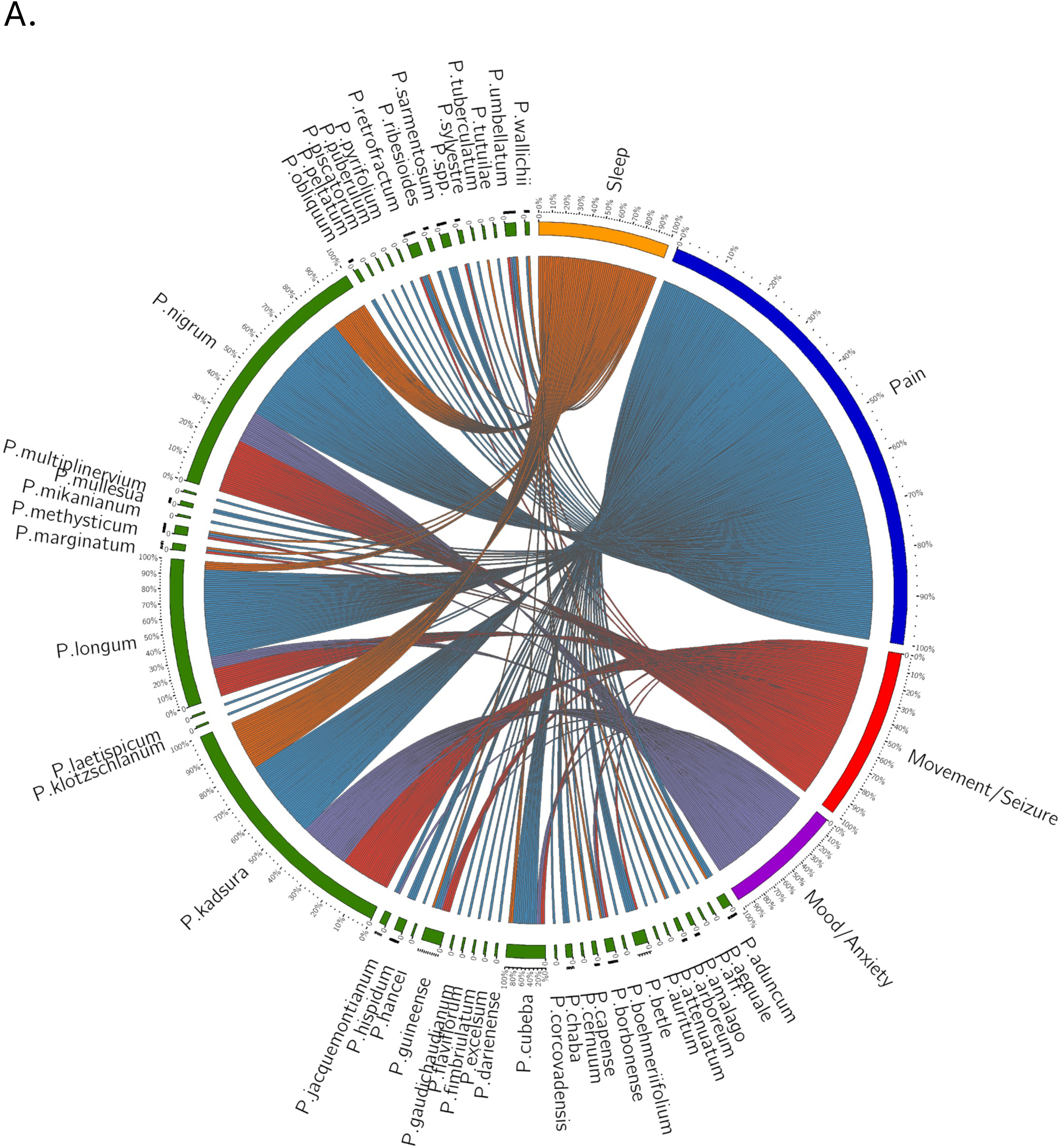
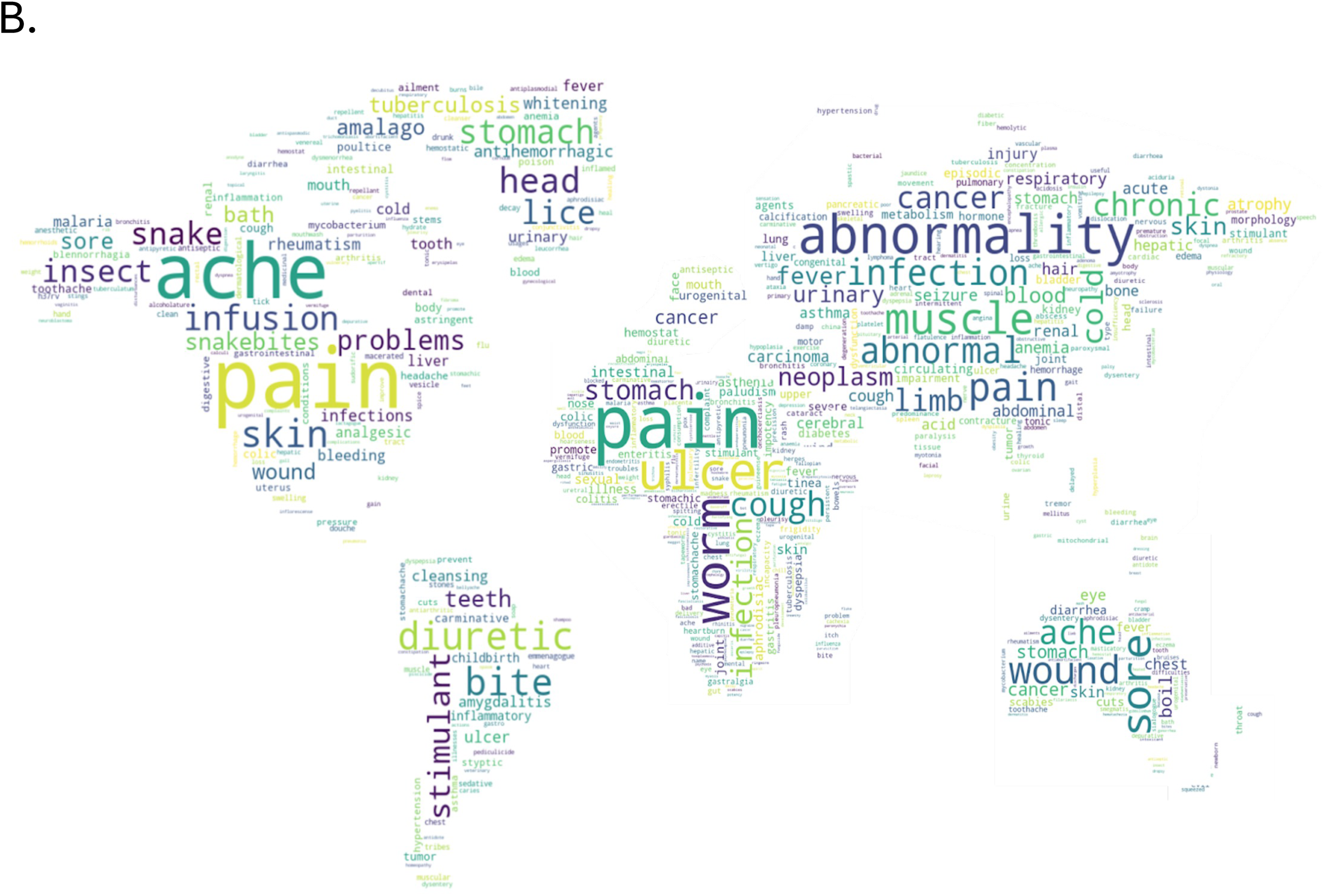
Diversity and global patterns of medicinal use for *Piper* species. **A. Circos plot of *Piper* species associations with major therapeutic indication categories across global traditional medical systems.** This visualization illustrates the frequency-based relationships between individual *Piper* species and four key therapeutic categories: Pain (blue), Movement/Seizures (red), Anxiety/Mood (purple), and Sleep (orange). Each connection line represents a documented instance where a *Piper* species is used in a traditional medicinal formula associated with one of these indications. Multiple lines between a species and different indication categories reflect formulas used to treat more than one disorder. Likewise, when multiple *Piper* species appear in the same multi-indication formula, each pairing is represented individually (e.g., a formula using *P. longum* and *P. nigrum* to treat all four categories results in total linkages). Notably, *Piper nigrum*, *Piper longum*, and *Piper kadsura* are most frequently associated with formulas targeting these indications. *Piper methysticum* (Kava) also shows strong multi-indication representation. Eight species (including *P. methysticum*, *P. borbonense*, *P. cubeba*, *P. guineense*, *P. kadsura*, *P. longum*, *P. nigrum*, and *P. umbellatum*) are documented in treatments across all four categories, with an additional nine species showing associations with at least two indication domains. These data support the hypothesis that therapeutic potential within the *Piper* genus extends well beyond *P. methysticum*, offering a broad foundation for polypharmaceutical or alternative formulations in neuropsychiatric and pain-related disorders. **B. Regional variation in medicinal indication patterns of *Piper* species use across global integrative medical systems.** Geospatial summary of *Piper* genus use cases across five GBIF regions (Americas, Africa, Europe, Asia, and Oceania) based on aggregated ethnomedical data. The word cloud is overlaid onto a world map, with each region displaying terms corresponding to medical indications associated with local *Piper* species and word size reflecting frequency with which a given indication appears in regional formulations.

As an alternative visualization, Figure 3B summarizes the geographical distribution of indications for *Piper. spp* across GIMS, excluding *Piper methysticum*. This figure illustrates the global distribution of Piper species use cases across five major geographical regions: the Americas, Africa, Europe, Asia, and Oceania. A word cloud is overlaid on each region of the world map, depicting the various medical indications treated by a species in the *Piper* genus in that specific region. The size of each word corresponds to the number of local formulas used to treat the given indication, highlighting the most common uses.

### Chemical composition and candidate compound analyses for efficacy in Mood/Anxiety, Sleep, Pain and Movement/Seizure indication categories in Piper methysticum and the wider Piper genus

One of the goals of this study is to evaluate, both in *Piper methysticum* and non KL-containing *Piper spp*, the potential for compounds other than KL to play a role in therapeutic efficacy for the 4 major indication categories of interest. We are interested in two related questions: Are there compounds other than KL that contribute to efficacy in *Piper methysticum*, and do they overlap with compounds that would be candidates for efficacy in *Piper spp.* that do not contain KL? In a first step we constructed a summative bioactive metabolome for *Piper methysticum*, drawing on multiple published studies that analyze the Piper methysticum phytochemistry, but using different extraction and characterization methodologies. This virtual metabolome is an important step to overcome the relative paucity of chemical data on *Piper methysticum* in available databases and the few papers detailing the chemical composition of *Piper methysticum* outside of its kavalactones and chalcones. Figure 4A and B shows a survey of bioactive chemical representation in *Piper methysticum* where we consolidated multiple data sources along with current literature on *Piper methysticum*. Figure 4A assembles a ‘meta-metabolome’ for Piper methysticum aggregating data form published extraction studies Parmar [73] and Salehi [74] are the most extensive literature compilations of chemical constituents for each species in the Piper genus. Xuan [75], Jaiswal [72], and Cheung [76] are the most extensive extraction papers on *Piper methysticum*. MESH (2017) [77] was the only traditional medicine database compiled for this study that listed chemical constituents for *Piper methysticum*. Pubchem [78] and and Dr. Duke’s Phytochemicals [37,38] contributed additional compounds and alternative forms of known *Piper methysticum* compounds to the list. Collectively among the presented sources, there are 269 recorded *Piper methysticum* compounds, of which are 268 recorded *Pm* compounds, of which there are 29 alkaloids, 31 benzenoids, 39 terpenes, 28 cinnamic acids, 10 flavonoids, 35 kavalactones, 17 chalcones, and 79 miscellaneous compounds. Figure 4B reconciles the chemical constituents with extraction methodology. While a definitive list of constituents has yet to be determined and this is largely influenced by naming conventions, extraction procedures, and plant variations, we were able to compile a composite ‘meta-metabolome’ for use as a starting point for linkage analysis with indications and comparisons with non-*Piper methysticum spp*.

**Figure 4.**
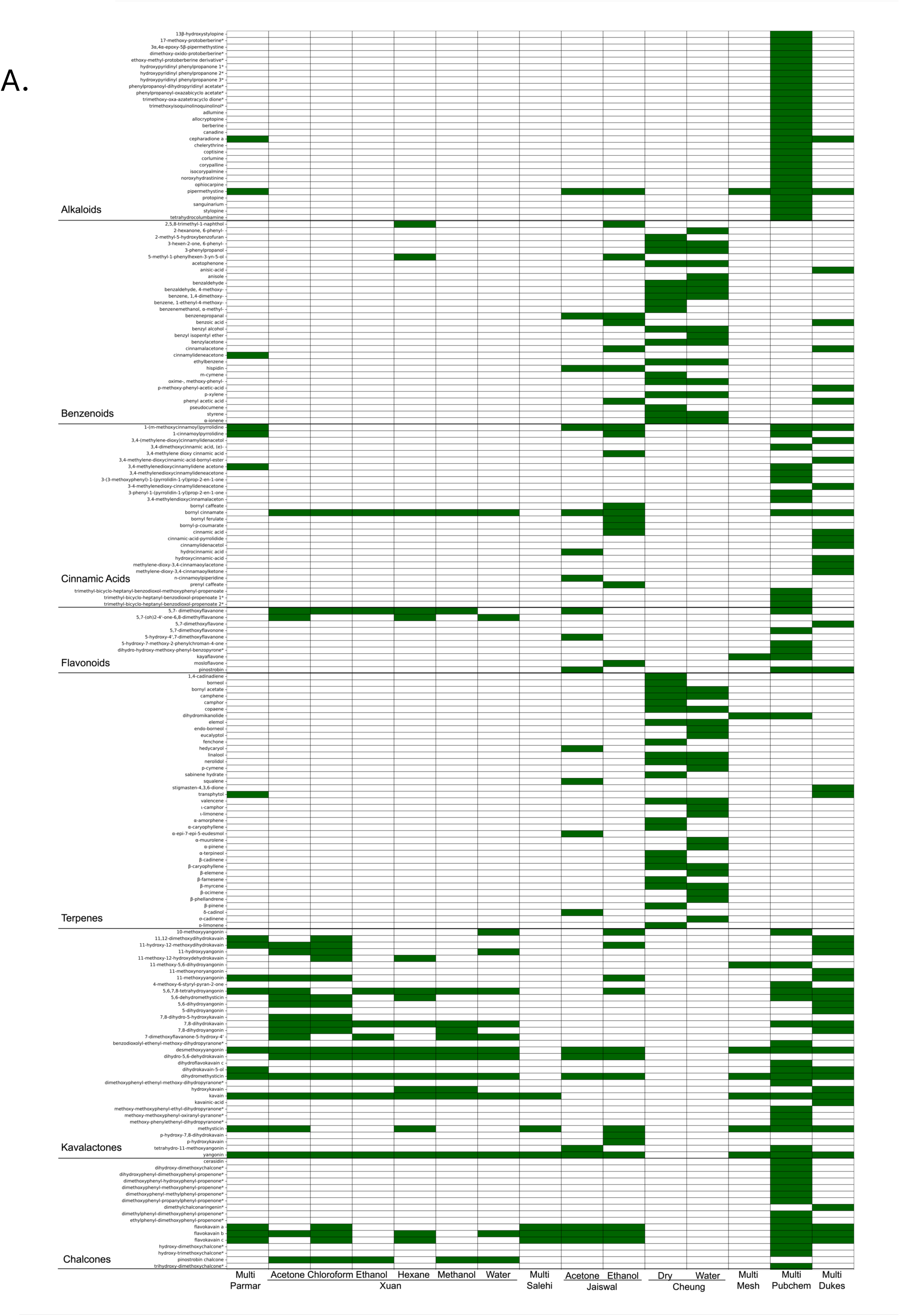
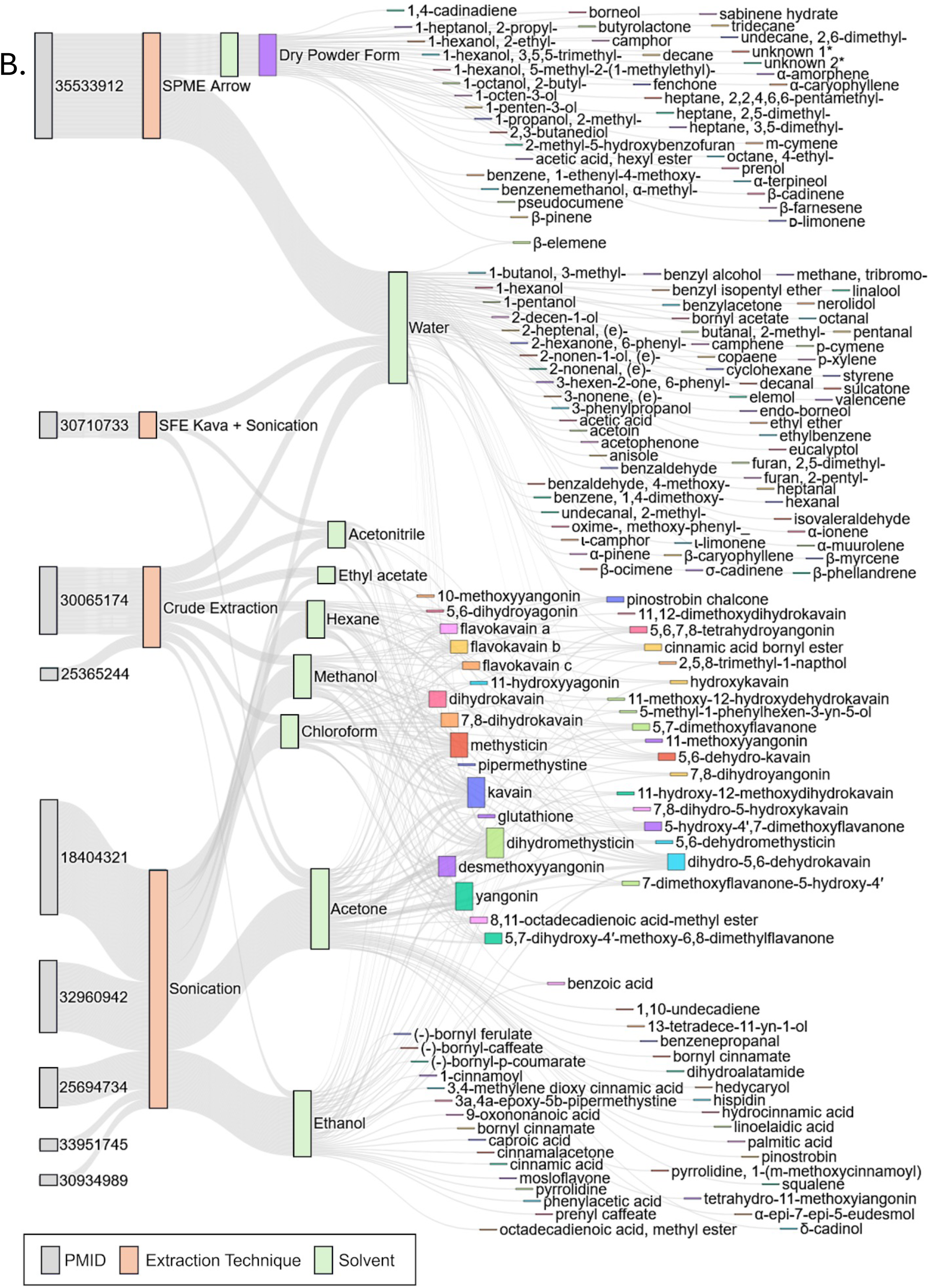
Chemical diversity and extraction methodology of *Piper methysticum*. **A. Composite heatmap of bioactive chemical constituents identified in *Piper methysticum* across data sources and extraction solvents.** This heatmap presents a curated summary of 268 secondary metabolites reported in *Piper methysticum* (Piper methysticum), categorized by major phytochemical classes: alkaloids, benzenoids, cinnamic acids, flavonoids, terpenes, kavalactones, chalcones, and miscellaneous compounds. Each row corresponds to a unique compound, while columns represent literature reviews, chemical databases, and extraction studies grouped by solvent system (e.g., acetone, ethanol, methanol, water). Green shading indicates the confirmed presence of a compound (either by successful extraction using the specified solvent in experimental studies or by inclusion in aggregated chemical databases and literature compilations). Key sources include Parmar (1997), Salehi (2019), Xuan (2008), Jaiswal (2020), Cheung (2022), MESH (2017), PubChem, and Duke’s Phytochemical Database. This provides an highlights methodological and solvent-dependent variation in compound detection and aggregates prior studies examining Piper methysticum constituents. **B. Sankey diagram linking published studies, extraction methods, solvents, and extracted compounds in *Piper methysticum*.** From left to right, the diagram connects: (1) PubMed IDs (PMIDS) of source studies (gray), (2) extraction techniques used (orange), (3) solvents employed (green), and (4) the specific compounds extracted (dark blue). The figure synthesizes data from major experimental studies utilizing methods such as solid-phase microextraction (SPME), sonication, supercritical fluid extraction (SFE), and crude solvent-based extraction. Solvents include water, ethanol, methanol, acetone, chloroform, hexane, ethyl acetate, and acetonitrile.

The bubble heatmap in Figure 5 depicts the strength of association between known *Piper methysticum* compounds in the compiled ‘meta-metabolome’ shared with other plant species and formulations used for pain, seizure/movement, anxiety disorders, and sleep disorders. The size of the bubbles represents the number (n) of medicinal formulas containing each compound and the heatmap colors indicate the percentage of these formulas treating a specific indication. The analysis reveals that for the majority of non-kavalactone compounds shared between *Piper methysticum* and alternative source Piper plants, there is definite association with the studied indications of anxiety/mood, seizure, sleep, and pain. This finding underscores the potential central role of kavalactones in inducing the observed medicinal effects. Noteworthy exceptions include canadine, anisole, camphor, and 1-cinnamoylpyrrolidine, which show associations with all four indications of interest. Figure 6 presents a network view, based on the assembled ‘meta-metabolome’ projecting the relative sharedness of compound sets between *Piper methysticum* and the non-*Piper methysticum* Piper species in PhAROS^TM^ that are associated with indications of interest in this study. This projection has some visual biases arising from the quantitative dominance of *Piper nigrum* in published metabolic studies, but it makes clear the specificity of association of canonical kavalactones with *Piper methysticum*. Figure 6 also highlights the differences between other *Piper spp.* in terms of overlapping and discrete metabolomes, and relative presence of the major compound families (e.g., alkaloids, terpenes, phenylpropanoids).

**Figure 5.**
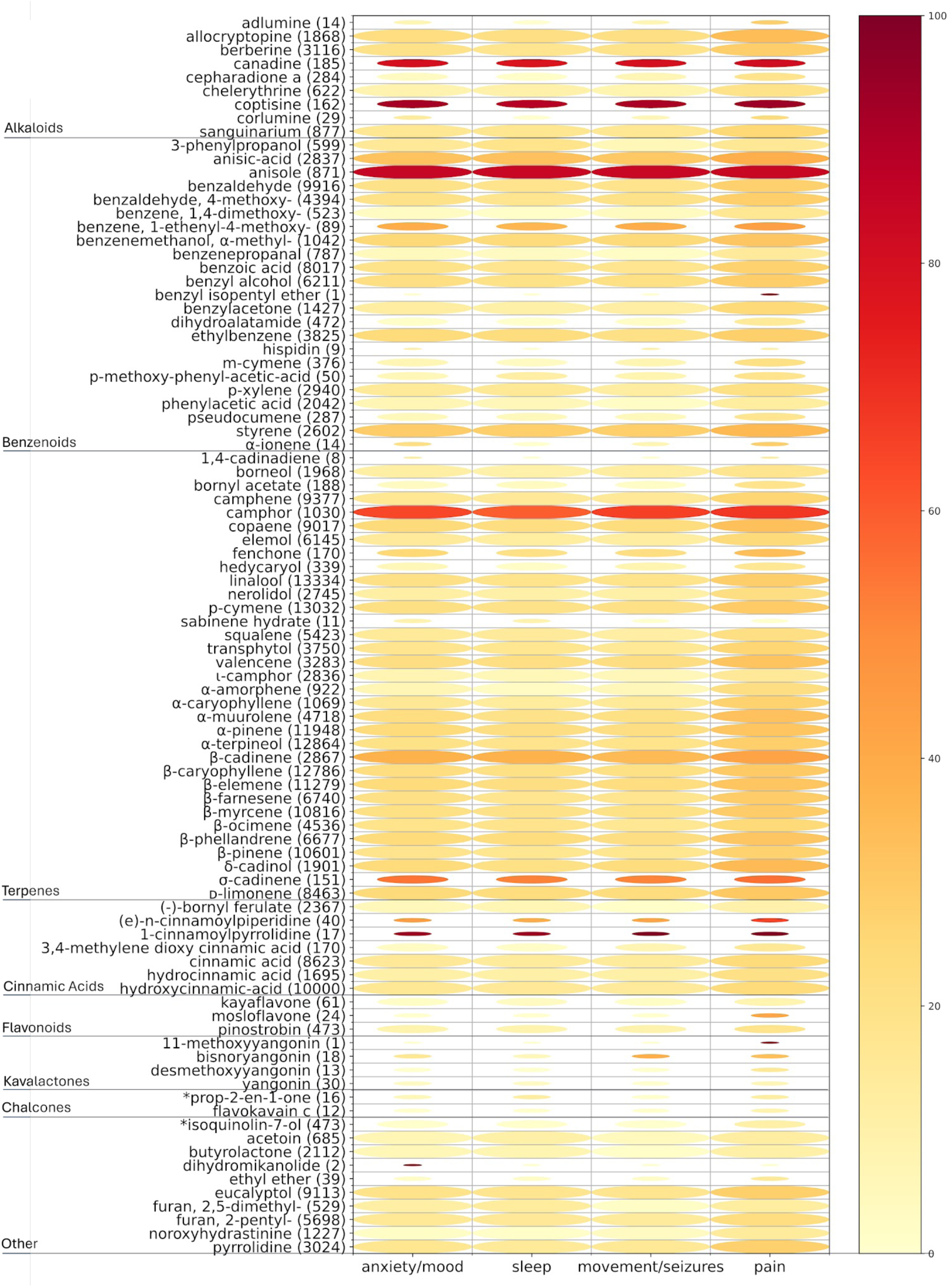
Bubble heatmap depicting indication-specific associations of *Piper methysticum* compounds shared with other plant species. This visualization assesses the strength and breadth of therapeutic associations for known *Piper methysticum* compounds that are also present in other plant species and medicinal formulations. Rows correspond to unique compounds, grouped by major chemical classes (e.g., alkaloids, terpenoids, cinnamic acids, flavonoids, kavalactones, chalcones). Columns represent four therapeutic indication categories of interest: anxiety/mood, sleep, movement/seizures, and pain. Bubble size reflects the number of traditional medicinal formulations (n) in which a given compound appears. Color intensity corresponds to the percentage of those formulations used to treat the specific indication, with darker shades indicating higher relevance. Most non-kavalactone compounds exhibit relatively limited association with the four therapeutic domains, a subset (e.g., anadine, coptisine, anisole, camphor, and 1-cinnamoylpyrrolidine) show cross-indication presences across all categories, suggesting that certain non-kavalactone constituents may contribute broadly to therapeutic outcomes. Some compounds have marked apparent specificity within the four indications of interest (e.g., some yangonin species showing preference for association with movement and pain).

**Figure 6.**
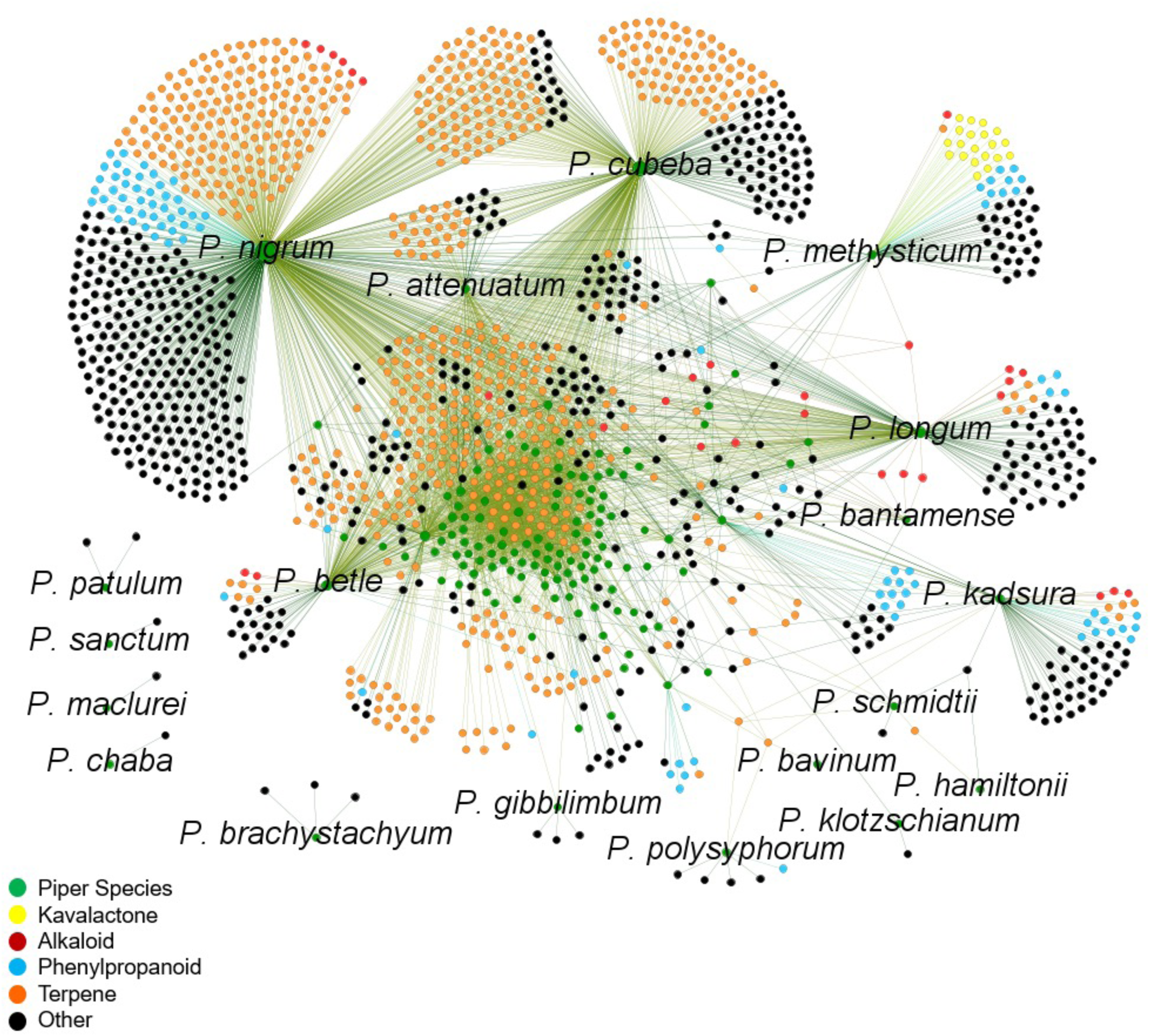
Network map of *Piper* species and shared chemical constituents across the genus. This network visualization displays chemical connectivity across 139 *Piper* species, based on shared secondary metabolites compiled from phytochemical datasets. Green nodes represent individual *Piper* species, while all other nodes represent distinct compound types: kavalactones (yellow), alkaloids (red), phenylpropanoids (cyan), terpenes (orange), and other compounds (black). Edges represent the presence of a given compound within a particular species. The map reveals clustering of chemically dense species such as *P. nigrum*, *P. methysticum*, *P. longum*, and *P. cubeba*, with partially overlapping chemical profiles. Notably, canonical kavalactones cluster exclusively with *P. methysticum*, emphasizing their species-specific presence while other pharmacologically relevant compound classes (e.g., alkaloids and terpenes) show broader distribution across the genus.

### Medicinal indications analysis of non-Piper methysticum Piper compounds and wider ingredient organism representation

Looking outside the *Piper methysticum*-specific kavalactones, we asked how compounds that are present in non-Piper methysticum *Piper spp.* associate with indications of interest in this study. Figure 7 shows a heat map/bubble plot visualization of potential alternatives to kavalactones that are present in *Piper spp.* other than *Piper methysticum*, and that are used in formulations that treat anxiety/mood, sleep, movement/seizures and pain. This is a first-pass candidate list for further study as therapeutically-active alternatives to KL. We note that some of these compounds are specific to certain non-*Piper methysticum* species (e.g. Kadsurenin) but that some are found in *Piper methysticum* also (e.g. cinnamic acid), suggesting potential therapeutic roles of compounds outside the canonical KL even when the therapeutic is a *Piper methysticum* extract.

**Figure 7.**
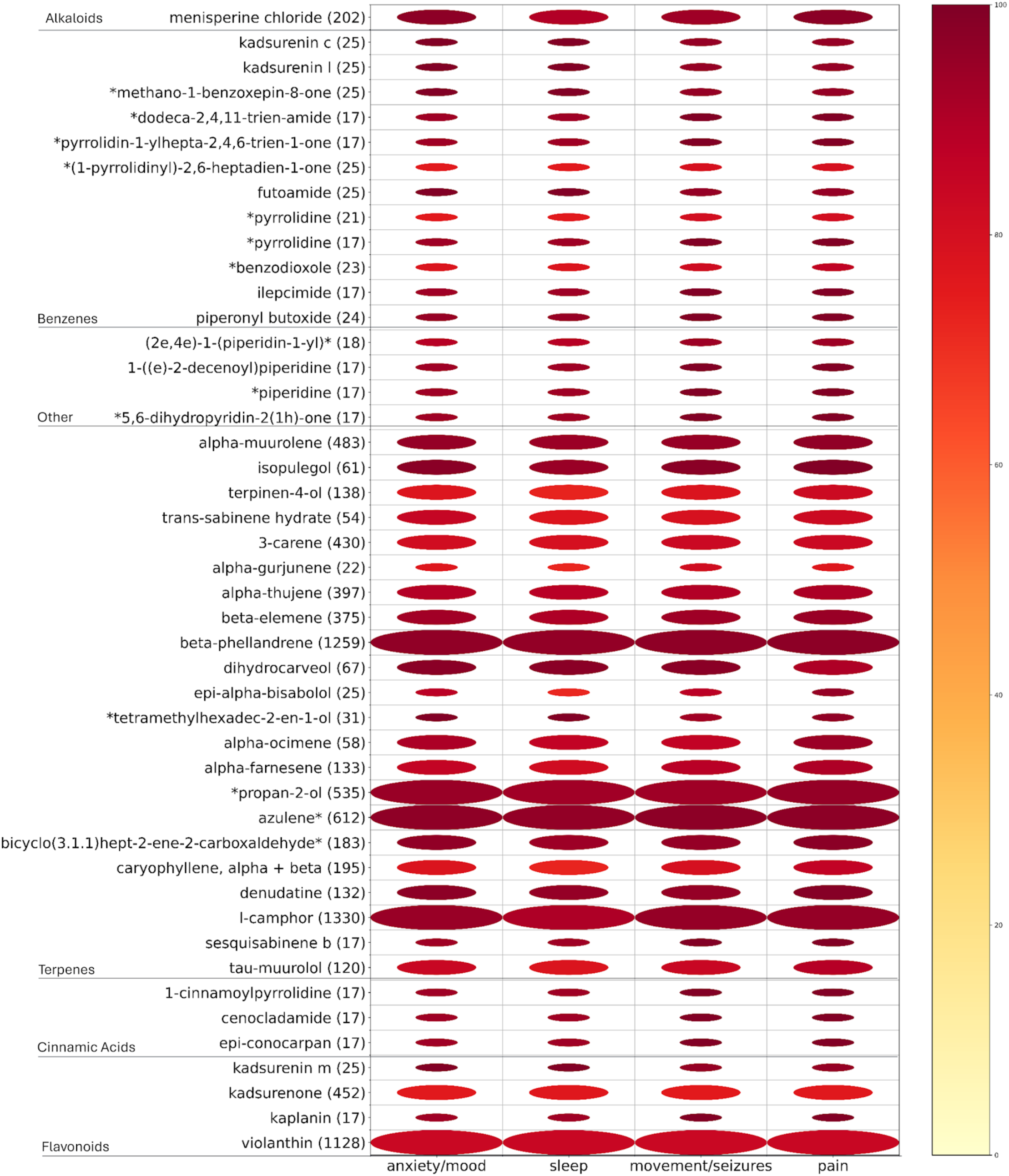
Medicinal indication analysis of non-*Piper methysticum* compounds: potential alternatives to kavalactones. Heatmap visualizes associations between bioactive compounds found in *Piper* species other than *P. methysticum* and their documented use in treating anxiety/mood disorders, sleep disturbances, movement/seizure-related disorders, and pain. Rows represent a compound (grouped by chemical class), columns represent indication categories. Bubble size corresponds to the number of traditional medicinal formulations in which a compound appears, while color intensity reflects the proportion of those formulations used to treat a given indication. Of 1162 Piper compounds associated with the indications of interest, 74 (shown here) had a sum of indication-specific percentages exceeding 300.Certain non-kavalactone candidates including terpenes (e.g., beta-elemene, beta-phellandrene), cinnamic acid derivatives, alkaloids (e.g., menisperine chloride), and flavonoids (e.g., violanthin) show strong multi-indication associations. These findings suggest chemically diverse alternatives to kavalactones that may underlie the efficacy of *Piper* species used outside the Pacific for these indications.

Figure 8 highlights the potential of exploring non-*Piper spp* for possible alternatives or combinatorial approaches to formulations addressing the indication of interest. For alkaloids, phenylypropanoids and terpenes (8A-8C) we evaluated the number of plant sources (ingredient organisms) in PhAROS™ as a source associated with that compound (upper panels). The compounds depicted represent those with the highest cumulative percentage across all four therapeutic categories of interest in this study (pain, anxiety/mood disorders, sleep disorders, and movement/seizure-related disorders) based on their use in phytomedicinal formulas.

**Figure 8.**
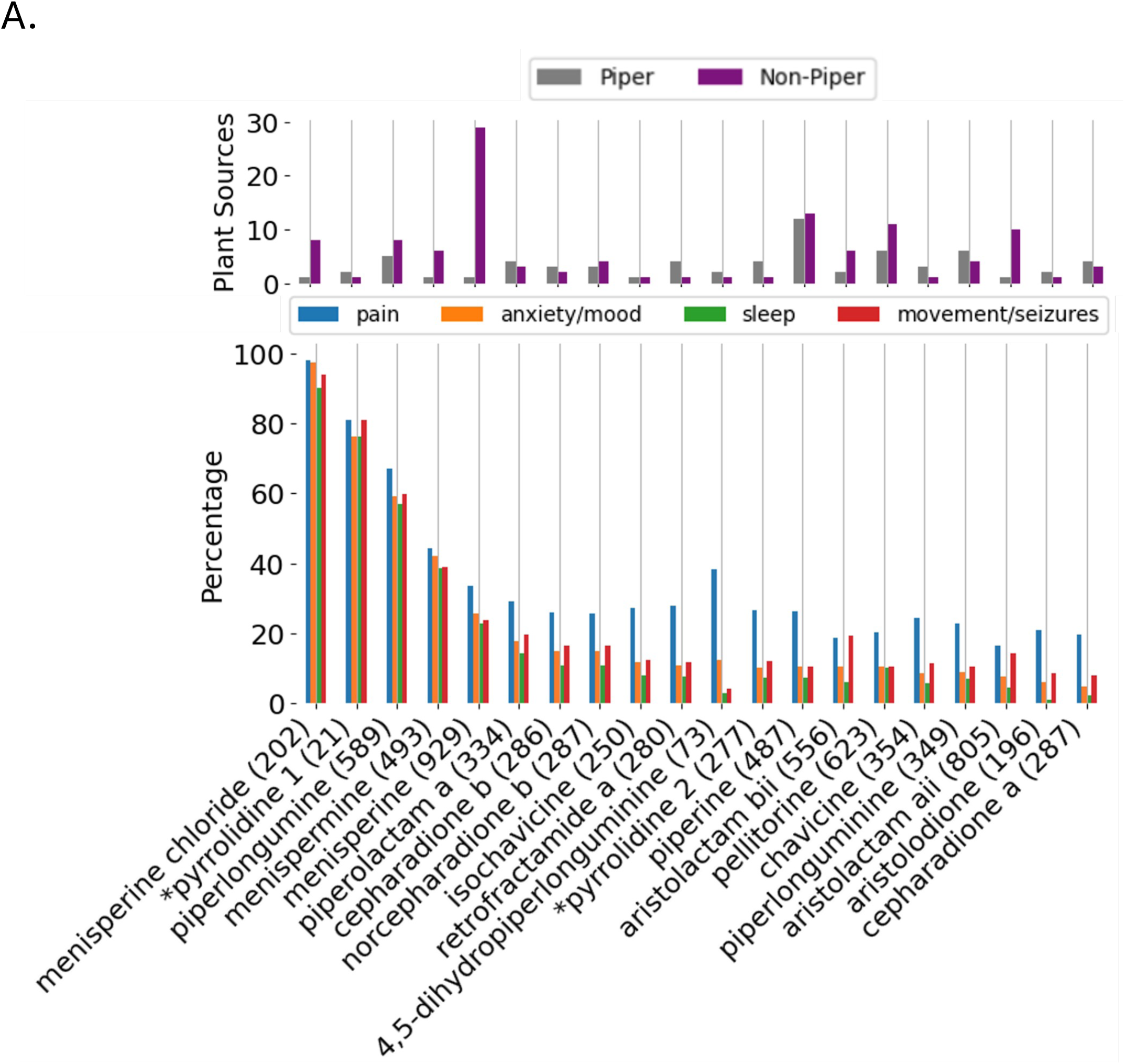
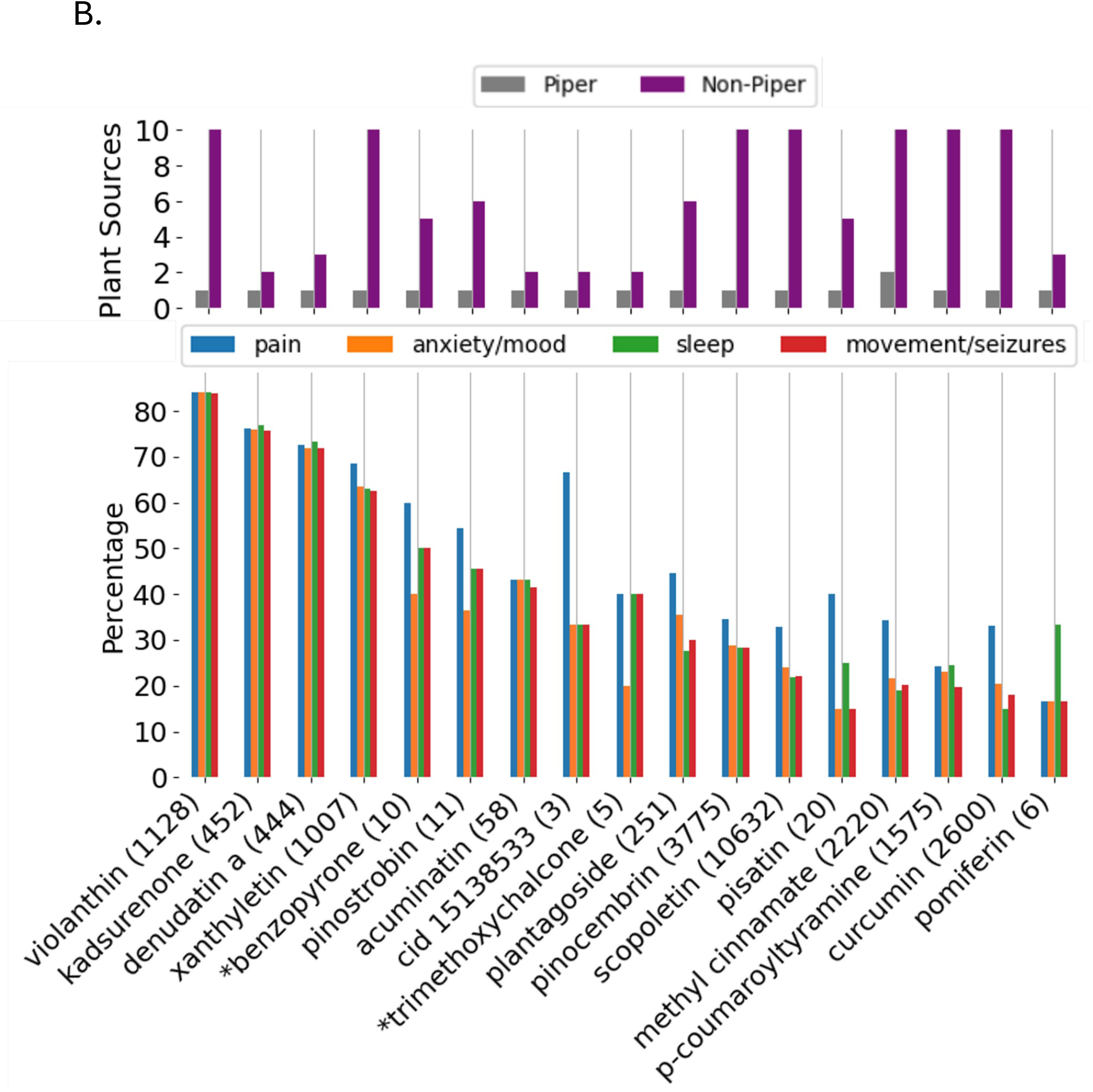
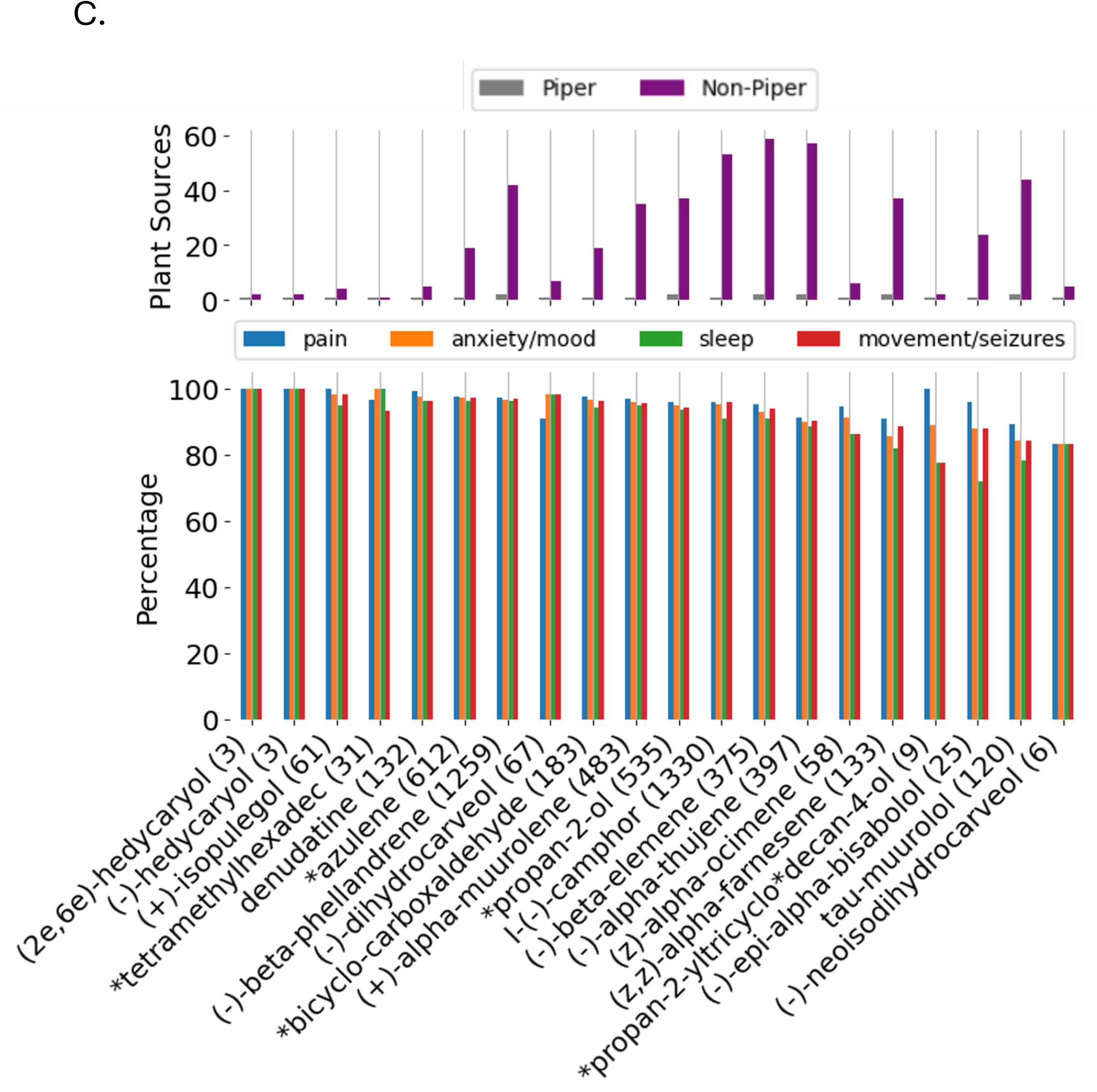
Therapeutic indication profiles and distribution of key phytochemicals within and beyond the *Piper* genus. **A. Piper alkaloids: medicinal indications and cross-species distribution.** *Lower panel.* Percentage of formulations containing each alkaloid used to treat pain (blue), anxiety/mood disorders (orange), sleep disorders (green), or movement/seizure-related disorders (red). *Upper panel.* Number of plant species that contain each alkaloid (gray = Piper, purple = non-Piper). Menisperine chloride, for example, appears in over 200 formulations, with >80% associated with one or more of the four indication categories. All alkaloids included in this analysis are present in at least one Piper species and one non-Piper species, suggesting pharmacological relevance and broad botanical distribution. *Pyrrolidine 1 = 1-[(2E,4E,8E)-9-(3,4-methylenedioxyphenyl)-2,4,8-nonatrienoyl]pyrrolidine. *Pyrrolidine 2 = 1-[(2E,8E)-9-(3,4-methylenedioxyphenyl)-2,8-nonadienoyl]pyrrolidine. **B, C. Visualization as in A, but for phenylpropanoids (B) and Terpenes (C**). **B.** Violanthin, kadsurenone, denudatin A, and xanthyletin exhibit particularly high association rates across all four indication domains. Several additional compounds such as *benzopyrone, acuminatin, and pinostrobin, also show moderate-to-strong linkage, underscoring the potential for broad-spectrum therapeutic utility beyond kavalactones. *Benzopyrone = (s)-2,3-dihydro-5-hydroxy-methoxy-2-phenyl-4-benzopyrone. *Trimethoxychalcone = 2’-hydroxy-4,4’,6’-trimethoxychalcone. **C.** *\*Tetramethylhexadec =* (E,7S,11R)-3,7,11,15-tetramethylhexadec-2-en-1-ol. *\*Azulene =* 1,2,3,4,5,6,7,8-octahydro-1,4-dimethyl-7-(1-methylethenyl)-, (1S,4S,7R). \**Propan-2-yltricyclodecan-4-ol* = (4r,10s)-4,10-dimethyl-7-propan-2-yltricyclo[4.4.0.01,5]decan-4-ol.

Figure 9 further underscores the potential of mining PhAROS™ for novel approaches to the indication of interest. In this visualization, we calculate a ‘compound index’ by identifying the percentage of all formulas for each indication investigated here (pain, anxiety/mood disorders, sleep disorders, or movement/seizure disorders) that contain the compound of interest and averaging the percentages across all four indications. The numbers of formulations with the compound of interest are then plotted on the x-axis, with the y-axis representing the compound index. Notably these data reflect only *Piper spp.* that have no documented association with the indications of interest based on PubMed searches.

**Figure 9.**
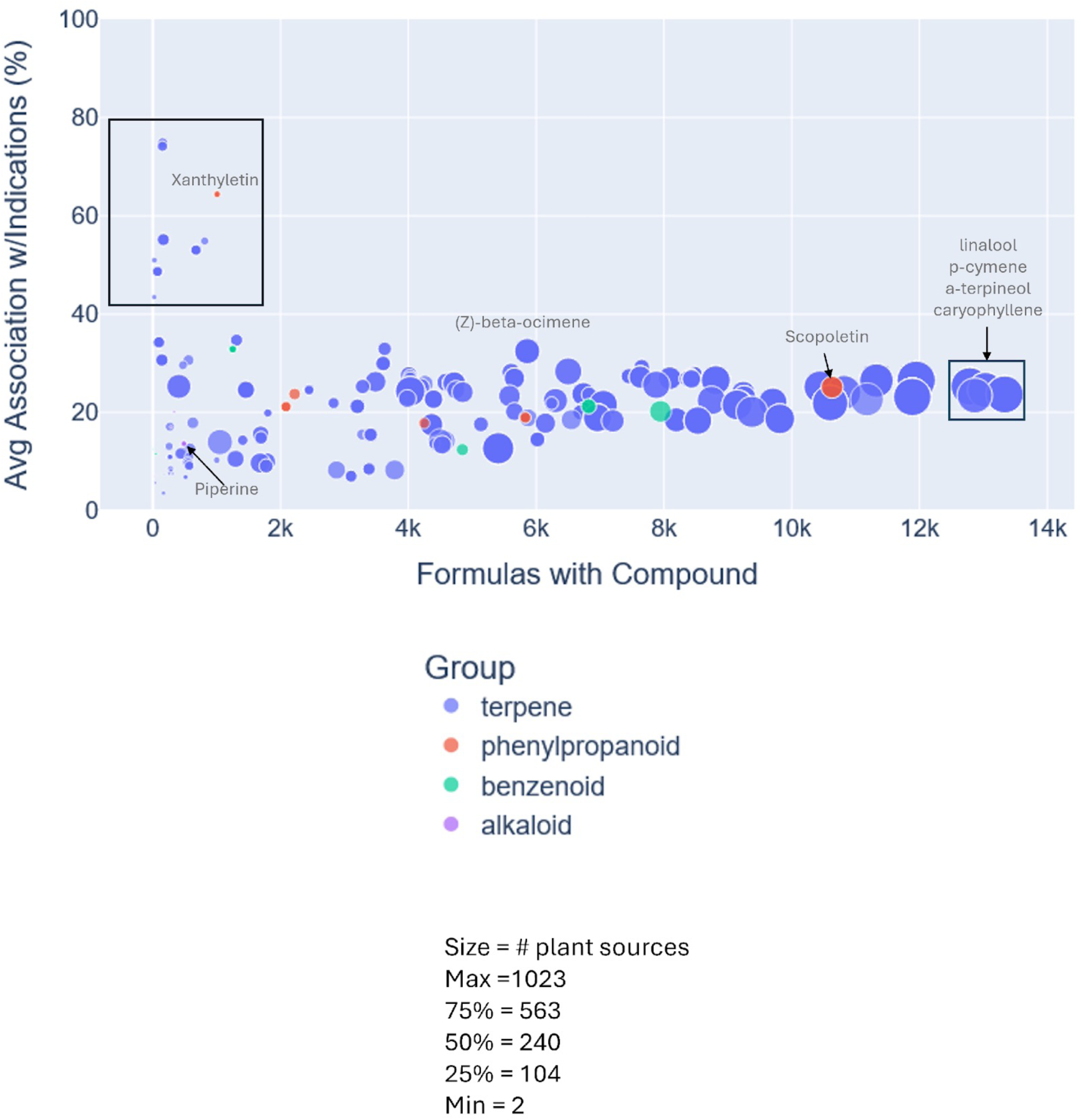
Chemical compound analysis of Piper Species with no PubMed associations to anxiety, pain, sleep or movement/seizure disorders. Scatter plot of compounds derived from 25 *Piper* species that, while GIMS-associated with treatment of anxiety, pain, sleep or movement/seizure disorders, have not been reported as connected to these indications in PubMed-indexed literature. Plots show number of formulations containing each compound by the average percentage of those formulas associated with the four therapeutic domains of interest. Dot size corresponds to the number of unique plant sources per compound, colored by chemical class: terpene (blue), phenylpropanoid (red), benzenoid (green), and alkaloid (purple). Highlighted compounds include Terpenes such as linalool, p-cymene, α-terpineol, and caryophyllene, each found in over 12,000 formulations and demonstrating substantial average association with the target indications, and Alkaloids such as piperine, piperolactam A, piperolactam C, and norcepharadione B, which vary widely in prevalence and therapeutic linkage. Several compounds (e.g., such as xanthyletin (a phenylpropanoid) and (Z)-β-ocimene (a terpene)) show high association with indications despite appearing in relatively few formulas, indicating strong but underrecognized therapeutic potential while *P. attenuatum*, *P. mullesua*, *P. obliquum*, and *P. hancei* are notable for contributing > 20 relevant compounds. This analysis identifies priority compounds and species for future investigation from underexplored *Piper* taxa with culturally validated but scientifically undocumented uses.

Decision support: further evaluation of *Piper methysticum/Piper spp*. candidates in *in vivo* model and druggability analyses.

As a final stage in this study, we piloted a decision support approach to assist in prioritizing compounds identified from *Piper methysticum* and the wider *Piper spp.* family for potential further evaluation in the indication of interest. In Figure 10 A-L we used a zebrafish (*Danio rario*) model system which has been validated for assessment of sedative and anti-anxiolytic properties in a medium throughput format. We used zebrafish larval behavior and thigmotaxis assays previously validated in anxiety [50]. We evaluated a subset of *Piper methysticum* compounds including KL and non-KL, and non-*Piper methysticum* species compounds that scores highly in PhaROS for association with anxiety and mood disorders. Notable findings include that a non-KL component (yangonin) of *Piper methysticum* is effective in this model as an anti-anxiolytic, with reduced stress in the absence of sedation which we view as a desirable effect profile. The three canonical KL tested perform in the stress and anxiety reduction parameters but tend towards inducing sedation. Piperine, widely found in *Piper spp* associated with the indication of interest, performs similarly to yangonin and dihyrdromethysticin with a desirable effect profile. These data allow us to demonstrate a decision support layer for ranking/prioritizing compounds for further study based on *in vivo* data.

**Figure 10:**
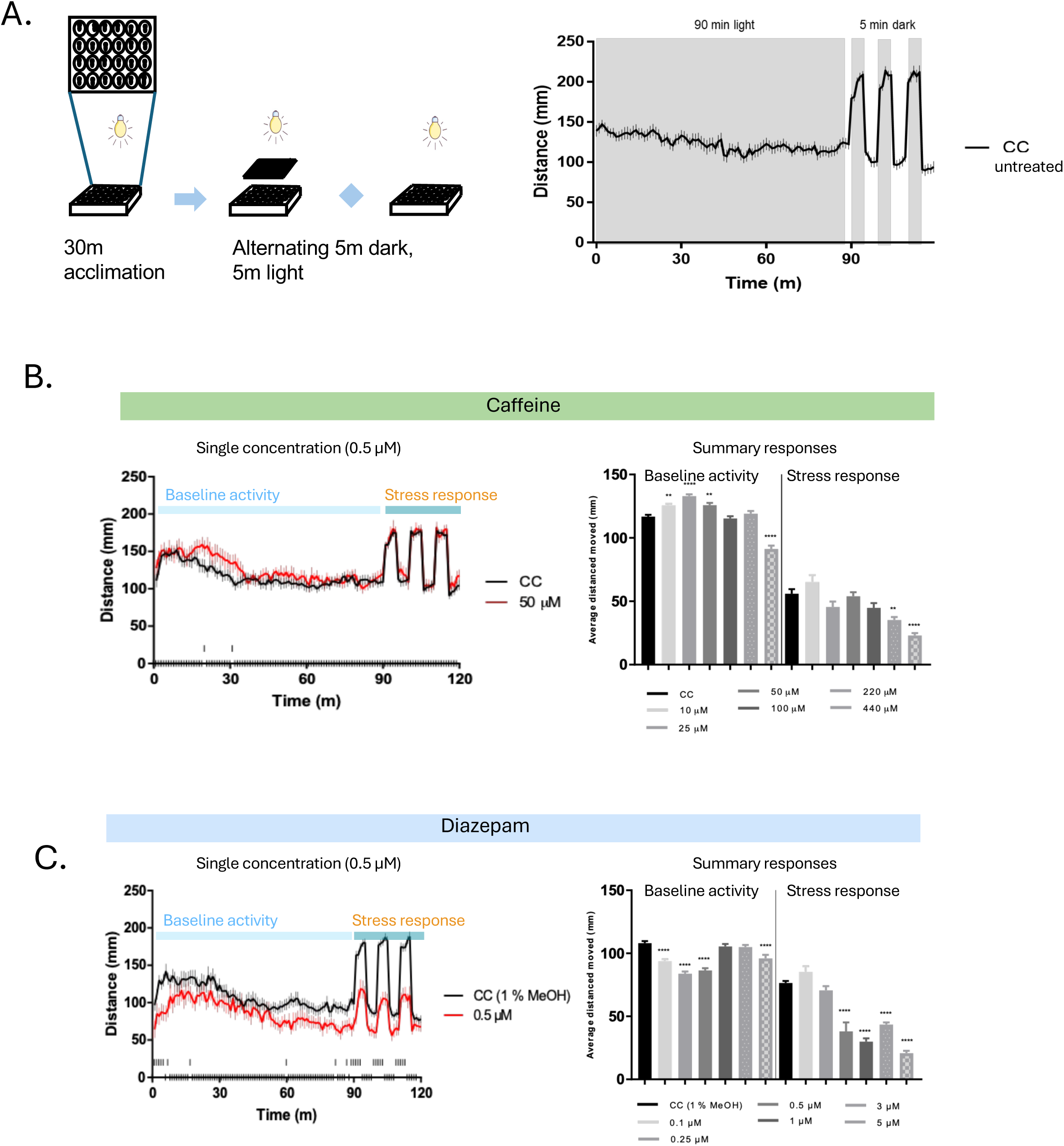
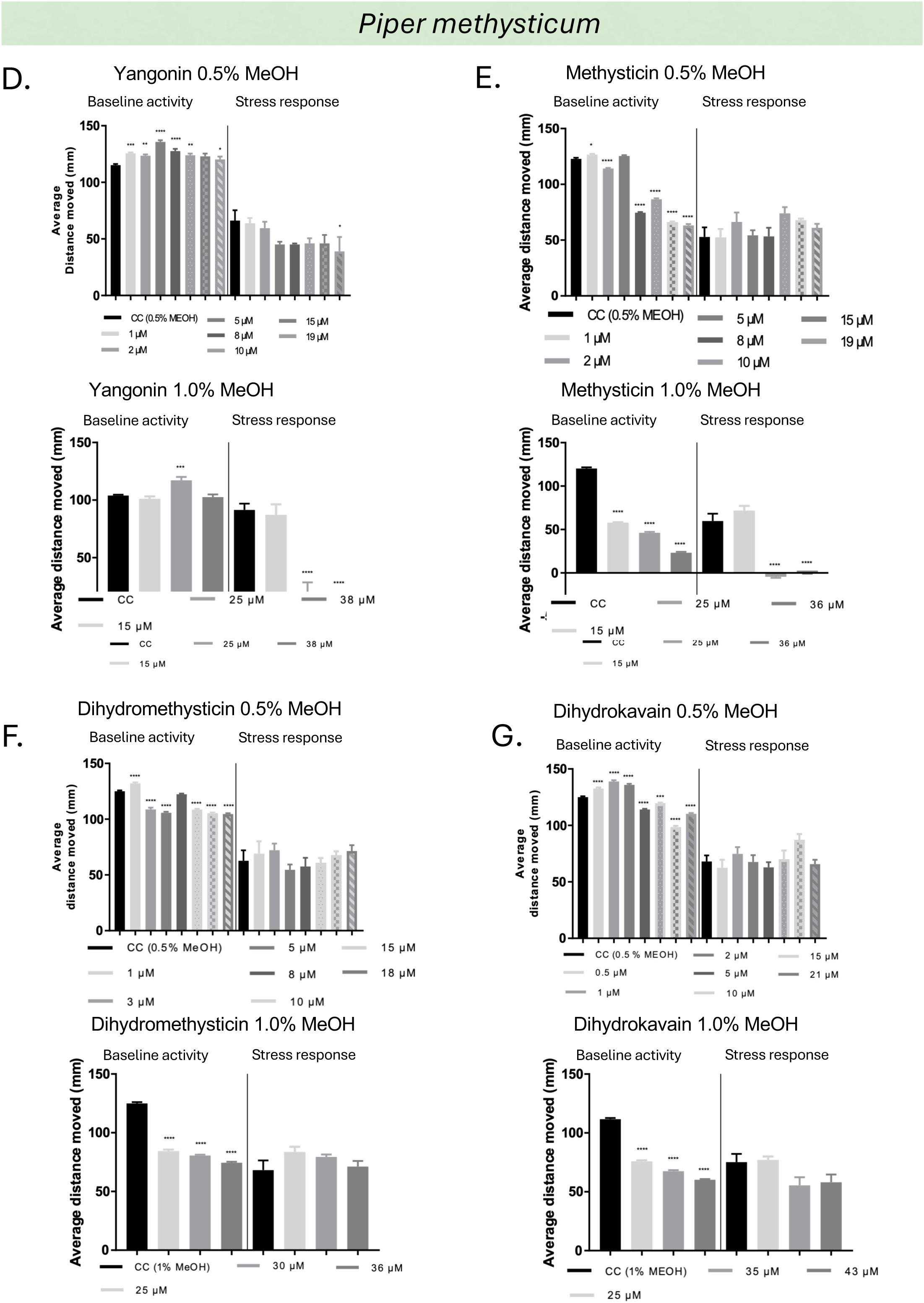
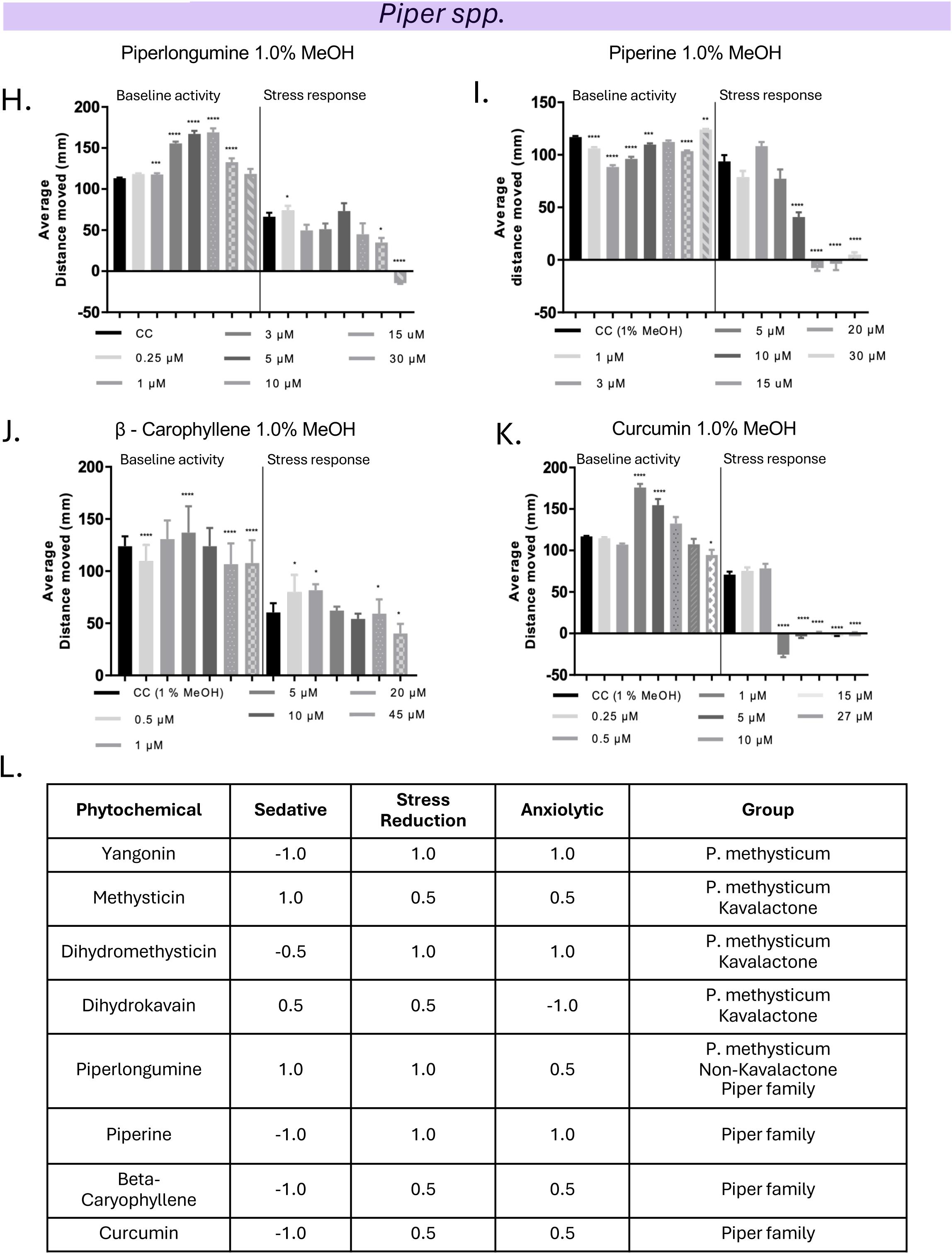
*Piper* family compounds have both sedative and anxiolytic effects on zebrafish larvae. **A.** Schematic representation of behavioral model and behavioral trace. Larvae are deposited into a 24-well plate and allowed to acclimate for 90 minutes in a light environment before an alternating light-dark stimulus is presented: 5 minutes of dark followed by 5 minutes of light. Increased activity in the dark is interpreted as flight behavior. Mean distance moved is presented in Summary Responses**, B-K.** Stress Response distance moved is determined by subtracting distance moved in 5 minutes light from distance moved in the following dark 5 minutes, averaged over 3 stimuli. **B.** Anxiogenic caffeine increases flight response at low concentrations but has little or sedating effects at high concentrations. **C.** Anxiolytic diazepam has broadly sedating effects at almost all concentrations. **D-G.** Summary responses for individual compounds derived from *Piper methysticum* solubilized in 0.5% methanol (top) or 1.0% methanol (bottom). **H-K.** Summary responses for individual compounds derived from medicinal plants in the *Piper* family solubilized in 1.0% methanol. **L.** Chart summary of figure 10 data.

PhAROS also has the capacity to investigate druggability indicators as a component of decision support workflows for candidate prioritization. Figure 11A, B and C illustrate NP-likeness, Lipinski Rule of 5 and wQED scores distributions compared to the entire PhAROS dataset for (i) the entire PhAROS™ data platform, (ii) the *Piper methysticum* virtual ‘meta-metabolome’ (Figure 4A), (ii) the non-*Piper methysticum Piper spp.* compounds associated with indications of interest (Figure 7) and finally the eight compounds tested in our *in vivo* pilot study (summarized in Figure 10L). We note significant enrichment of high-score druggability parameters in the various Piper compound sets. This provides discrimination potential as a component of the decision support goal of ranking and prioritizing compounds for inclusion in potential minimal essential therapeutic formulations (Figure 11).

**Figure 11.**
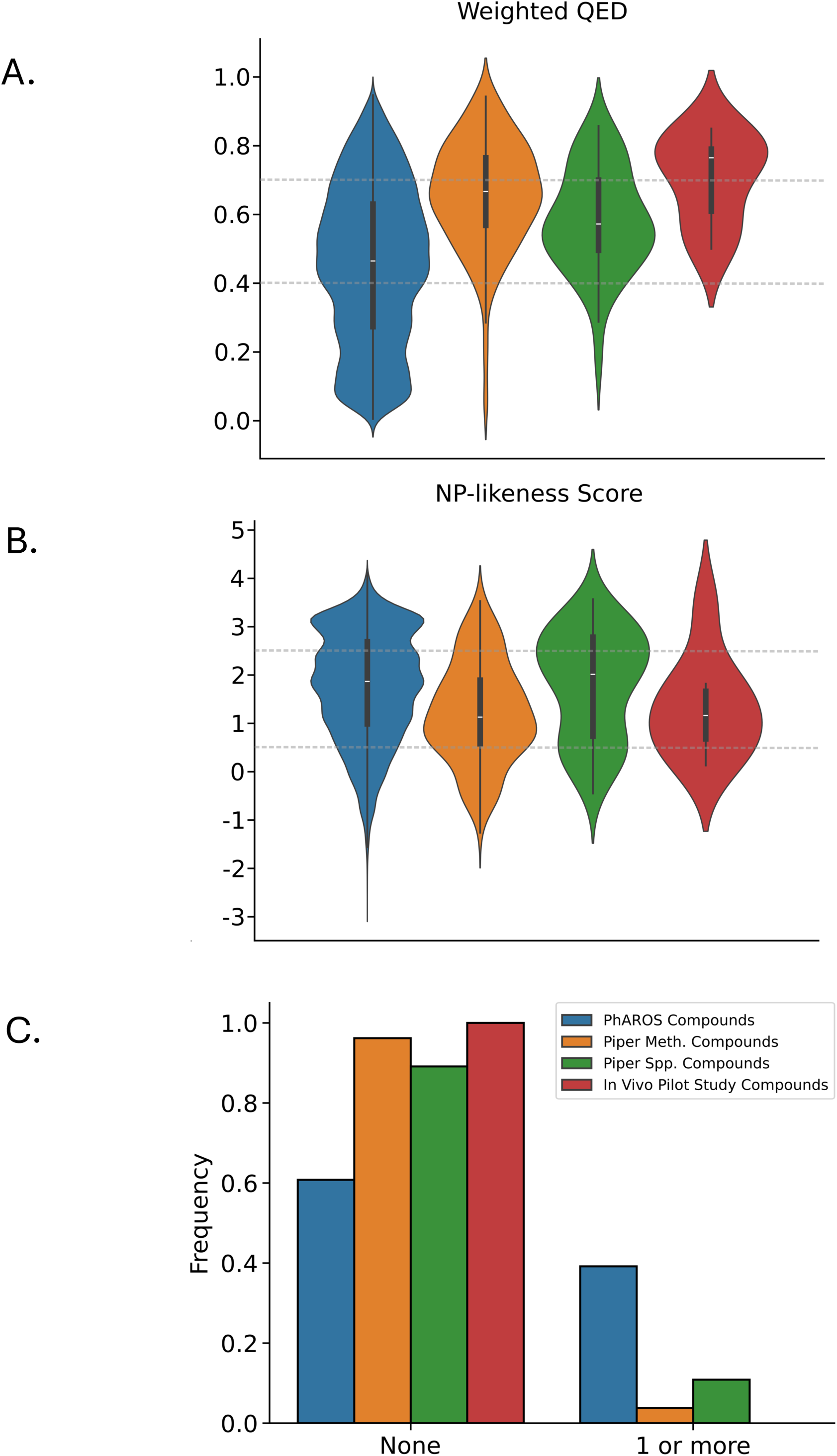
Visualization of druggability metrics for candidate compound groups. Visualizations of weighted quantitative estimate of drug-likeness (QED), natural product-likeness (NP-likeness), and the number of Lipinski’s Rule of Five (RO5) violations for four distinct sets of compounds: (1) all natural compounds contained in the PhAROS data platform (blue; left-most), (2) all natural compounds reported in the *Piper methysticum* secondary ‘meta-metabolome’ (orange; second from left), (3) all natural compounds found in other *Piper* spp. and associated with indications of interest (green; second from right), and (4) the eight natural compounds tested in our in vivo pilot study (red; right-most). All molecular properties were obtained using a combination of ChEMBL v32 database queries and the RDKit cheminformatics package (for compounds not found in ChEMBL). Note that these four sets of compounds are not mutually exclusive, and some compounds may appear in more than one set. **A.** Violin plot showing the distribution of weighted QED scores in the four sets of natural compounds. Dashed grey horizontal lines indicate the desirable range of weighted QED, from 0.4 to 0.7 (range: 0.0 to 1.0). **B.** Violin plot showing the distribution of NP-likeness scores in the four sets. Dashed grey horizontal lines indicate the desirable range, from 0.5 to 2.5 (range: –5 to +5). **C.** Grouped bar chart showing the proportion of compounds in each set with no Lipinski’s RO5 violations (left-most bars) versus those with one or more violations (right-most bars).

## Discussion

Sleep disturbances, anxiety disorders, pain, and seizure-related conditions represent ongoing areas of unmet clinical need, particularly where existing pharmacological treatments carry limitations related to efficacy, safety, or cultural relevance. This study investigated whether the therapeutic properties attributed to *Piper methysticum*, particularly in the treatment of anxiety, pain, sleep disturbances, and seizure-related disorders, originate exclusively from this species or extend across the broader *Piper* genus. Prior studies associate kavalactones with the anxiolytic effects of *Piper methysticum* [1–4]. However, we identified multiple non-methysticum Piper species, including *Piper nigrum*, *Piper longum*, and *Piper kadsura*, used traditionally in various global contexts for similar indications. These species lack KL but contain diverse classes of bioactive compounds such as alkaloids, phenylpropanoids, flavonoids, and terpenes, which our analysis linked to formulations treating anxiety, sleep disorders, pain, and seizures. We constructed multi-layered network visualizations and compound-association analyses that highlighted both overlap and divergence in the therapeutic chemistry among these species. These findings broaden the pharmacological relevance of the *Piper* genus and underscore the cultural and chemical plurality behind phytomedicine.

We developed and applied PhAROS^TM^, a computational platform that integrates chemical, ethnomedical, and biological data to identify, prioritize, and support decision-making around candidate compounds and species. Instead of relying on traditional bioassay-guided fractionation [79,80], which consumes significant time and material, we leveraged PhAROS™ to rapidly correlate plant-derived compounds with indications of interest using a structured query framework. This method enabled our team to extract meaningful relationships from across more than a dozen traditional medicine databases, covering diverse geographical and cultural origins. We enriched the compound metadata with identifiers, classification schemes, and chemical properties to assess not only presence but also relevance. PhAROS™ facilitated the comparison of canonical kavalactones with a spectrum of structurally and functionally diverse compounds found in both *P. methysticum* and other *Piper* species. These included piperine, canadine, kadsurenone, and cinnamic acid derivatives which are compounds with demonstrated links to neurological indications [81–85]. This platform reduces the complexity of traditional medicine formulations. making it feasible to construct prioritized candidate shortlists based on drug-likeness, formulation compatibility, and fidelity to the ethnomedical knowledge bases.

Our findings challenge the assumption that kavalactones solely mediate the therapeutic efficacy of kava [86]. Through compound-level analysis and cross-referencing with documented ethnomedical uses, we demonstrated that several non-kavalactone constituents, within *P. methysticum* and in other *Piper spp*., show high indication-specific association scores. Yangonin, for example, displayed anxiolytic-like activity without concurrent sedation in zebrafish behavioral assays, indicating a distinct and desirable pharmacodynamic profile and supporting prior studies that differentiate its effects from KL in other indications [87]. Piperine, common in *P. nigrum* and other widely distributed *Piper* species, exhibited a similar behavioral signature [81]. By identifying non-kavalactone candidates with therapeutic potential, we offer a pathway to reduce dependency on *P. methysticum*, a species with ecologically constrained cultivation and high cultural specificity. This redirection may also alleviate pressure on Pacific Island supply chains and mitigate the cultural and ethical concerns surrounding appropriation and commodification of culturally significant plants [88,89]. Our findings affirm the existence of culturally-validated, biogeographically distinct *Piper methysticum* alternatives that may support the same therapeutic goals.

Our work is contextualized in an ongoing debate on a range of *Piper methysticum* delivery modalities, from traditional kava usage to commercial (often chemical) extractions of whole plants or plant parts that are packaged as a nutraceutical. Pacific Island communities traditionally prepare kava as an aqueous infusion consumed within a relational, communal, and behavioral framework associated with *talanoa* [20]. Only within this context do users report the full spectrum of therapeutic effects, including anxiolysis, emotional catharsis, and restorative sleep. In contrast, commercially available extracts (which do not meet the cultural definition of kava but are better termed ‘products containing *Piper methysticum*’ (often produced using organic solvents) lack not only the broader phytochemical matrix but also the cultural structure that contributes to efficacy. These extracts exhibit altered chemical profiles, possibly concentrating pro-inflammatory compounds like flavokavains and excluding water-soluble protective components such as glutathione. As a result, products that neither maintain traditional composition nor achieve evidence-based optimization may pose elevated risks without delivering commensurate benefits. For practical translation across geographies and health systems, we propose two viable pathways: (1) recapitulate traditional formulations as closely as possible using naturalistic kava preparation only with chemical and behavioral components, or (2) construct new, culturally-informed multi-compound formulations that reflect traditional patterns of synergy and efficacy. These pathways de-emphasize the chemically aggressive, decontextualized preparations that distort both pharmacological integrity and cultural intent [52]. Our work suggest that it will be possible to rationally design ‘minimal essential effective formulations’ (MEEF) that recapitulate desired effects but reduce the complexity and attendant regulatory, consistency and supply issues that may attend working only with the whole *Piper methysticum* plant in naturalistic settings. We are also positioned to rationally design ‘transcultural’ MEEF that leverage inclusion of compounds outside *Piper methysticum*, from different medical traditions, but which have been identified for the same therapeutic goals. This ‘transcultural’ potential is a strength of PhAROS™ as the platform lets us assemble medicine that are de-siloed from geographical, cultural, linguistic and biodiversity boundaries.

In this context, PhAROS™ also supports a decision framework for formulation strategy by enabling the development of minimal essential combinations that preserve the core therapeutic mechanisms of naturalistic kava while reducing dependency on full-spectrum whole-plant extracts. By identifying key compounds (e.g., yangonin, piperine, and select phenylpropanoids) that appear consistently across efficacious preparations, formulators can design synergistic combinations that remain compatible with cultural/behavioral practices like *talanoa*. This strategy reduces stress on vulnerable kava supply chains, circumvents biogeographic limitations of *P. methysticum*, and offers a means to mitigate variability introduced by day-to-day or batch-to-batch differences in native plant material. Such formulations could serve as culturally respectful yet logistically feasible interventions that retain therapeutic authenticity while expanding accessibility across global health settings.

We encountered several limitations in the present study. Existing phytochemical profiles for many Piper species remain incomplete or inconsistent. Species like *Piper brachystachyum* and *Piper sanctum* have sparse chemical data, not necessarily because they lack activity, but due to limited research being performed on them. Variances in botanical nomenclature (vernacular, taxonomic, and synonymic) create additional challenges. Open source phytopharmacopeias can apply outdated or region-specific plant names, which complicated cross-dataset species mapping. Similarly, chemical identifiers varied significantly across sources, with inconsistencies in IUPAC names, PubChem CIDs, and structural formats such as SMILES or InChIKeys. These disparities hinder unambiguous compound matching. Although we mitigated some issues using canonical IDs and cross-referencing, we could not eliminate all issues related to data quality. We note also that our *in vivo* validation experiments covered only a subset of high-priority compounds, focusing primarily on anxiolytic behavior in zebrafish models. Ongoing and future work in this system is addressing pain, and seizure-related phenotypes, and starting to evaluate chronic exposure and multi-compound interactions (Small-Howard, MS in preparation).

Despite these limitations, the study provides a valuable proof-of-concept. It underscores the need for more exhaustive phytochemical characterization of medicinal plants and for greater standardization (or at least careful curation) of ethnobotanical data. Moving forward, we envision PhAROS^TM^ and its growing assets suite of computational tools supporting a new generation of culturally- and mechanistically-informed phytotherapeutics. In ongoing work, PhAROS^TM^ has been augmented with target and mechanism API and data layers, as well as CYP metabolism data. This provides the potential to elucidate the mechanisms (e.g. GABAergic, serotonergic, cannabinoid, or anti-inflammatory pathways in the case of *Piper methysticum* and *Piper spp*) through which the phytomedical formulations exert their effects and establish critical pharmacodynamic considerations related to side effects and personalized outcomes. This implies that we will be able to design rationally simplified interventions that respect the *multi-component, multi-target* nature of polypharmaceutical phytomedicine, potentially cross extant cultural and biogeographical boundaries and support accelerated development timelines, regulatory tractability and improved patient outcomes.

## Acknowledgements

This work has been supported by: GB Global Biopharma Research grant to HT. NIH AIM-AHEAD OT2OD032581-01 (AJS and HT). NIH INBRE P20 GM103466) (HT). NSF INCLUDES Alliance HRD-2217242 (HT, AJS). NSF RII Track I ESPCoR OIA-2149133 (HT, AJS). NIH 1R15DA051749-01 (HT). Health Research Council of New Zealand Pacific Project Grant, 25/153, to SA. The views and conclusions contained in this document are those of the authors and should not be interpreted as representing the official policies, either expressed or implied, of the NIH. The views and conclusions contained in this document are those of the authors and should not be interpreted as representing the official policies, either expressed or implied, of the NSF.

## Declaration of generative AI and AI-assisted technologies in the writing process

During the preparation of this work the author(s) used ChatGPT4.o for research and to edit for clarity. After using this tool the author(s) reviewed and edited the content as needed and take(s) full responsibility for the content of the published article.

## Author Contributions (CRediT Taxonomy)

Conceptualization: HT, AJS, ALSH. Data curation: BW, BR. Funding acquisition: HT, AJS, ALSH. Investigation: BR, BW, LE, JN. Methodology, BR, BW, LE, HT, ALSH. Software: BR, BW. Visualization, BR, BW, JN, AR. Writing – original draft: HT, CJ, CF, BR, BW, AJS, ALSH, JH. Writing – review & editing: HT, CJ, BR, BW, CNA, AR, AA, AJS, ALSH.

## Statement on use of Generative AI

Declaration of generative AI and AI-assisted technologies in the writing process during the preparation of this work the author(s) used iChatGPT4o in order to research, edit and check text. After using this tool/service, the author(s) reviewed and edited the content as needed and take(s) full responsibility for the content of the publication.

---

**Supplemental Figure A.**
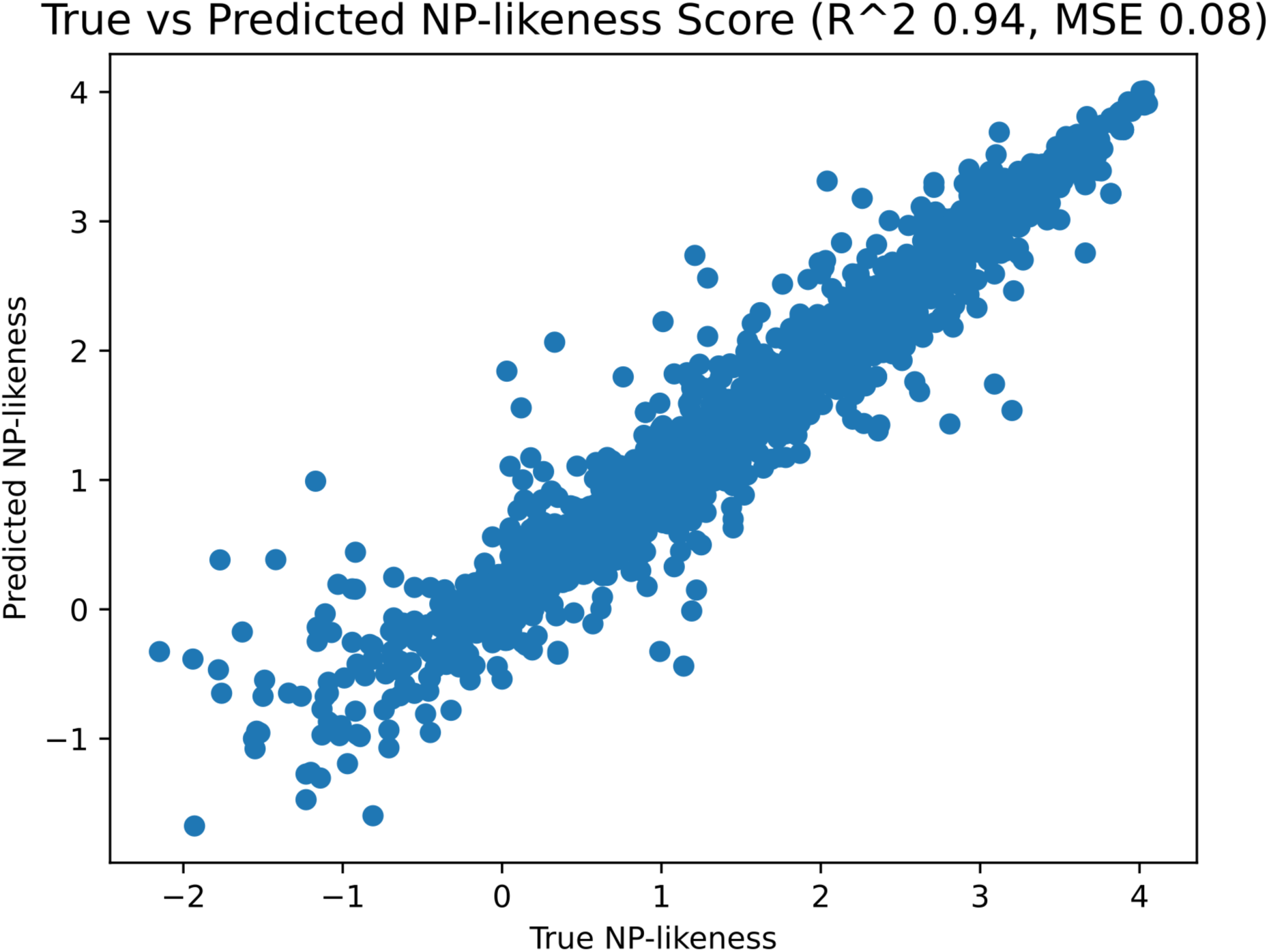
Assessment of true versus predicted NP-likeness score.

**Supplemental Figure B.**
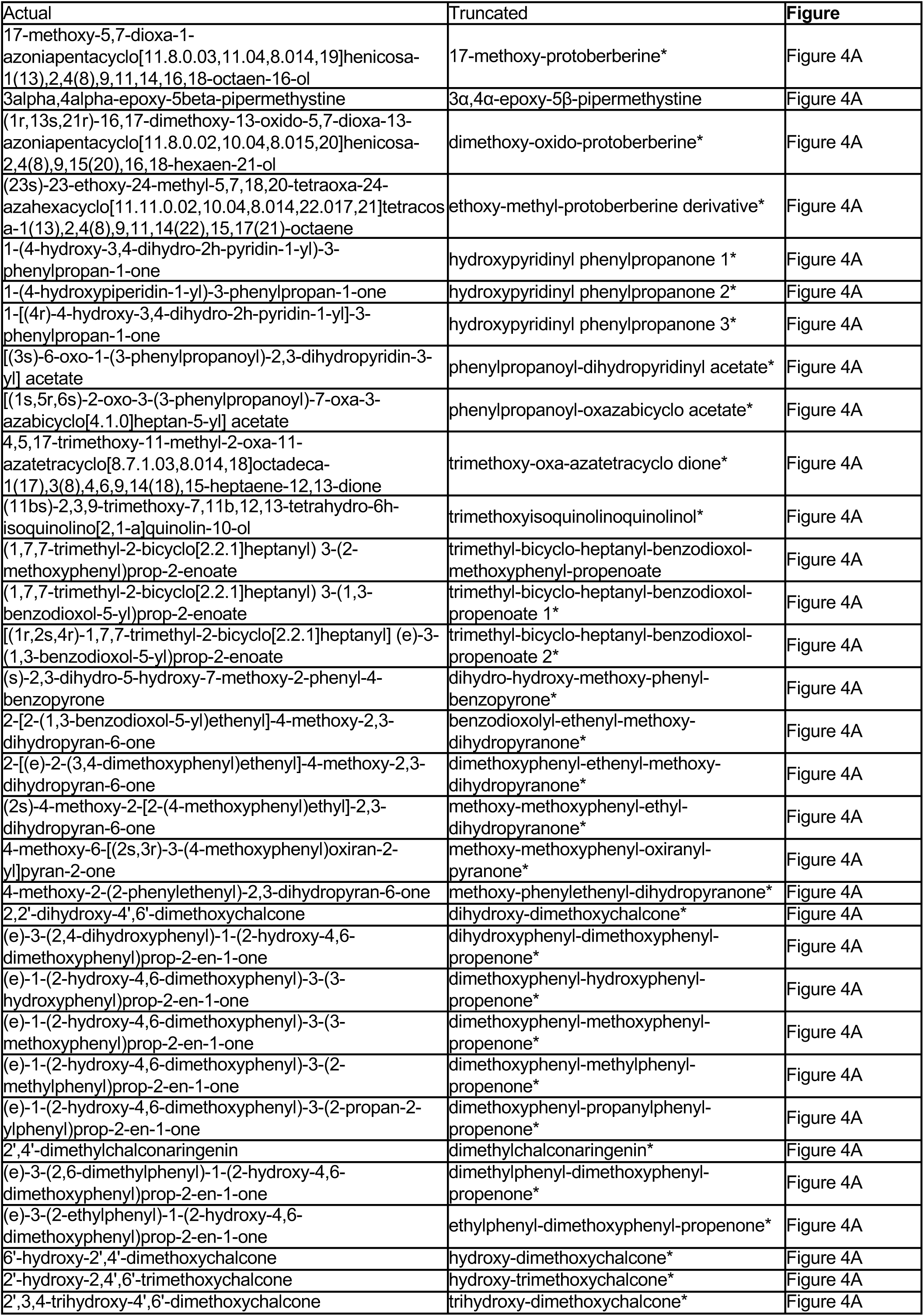
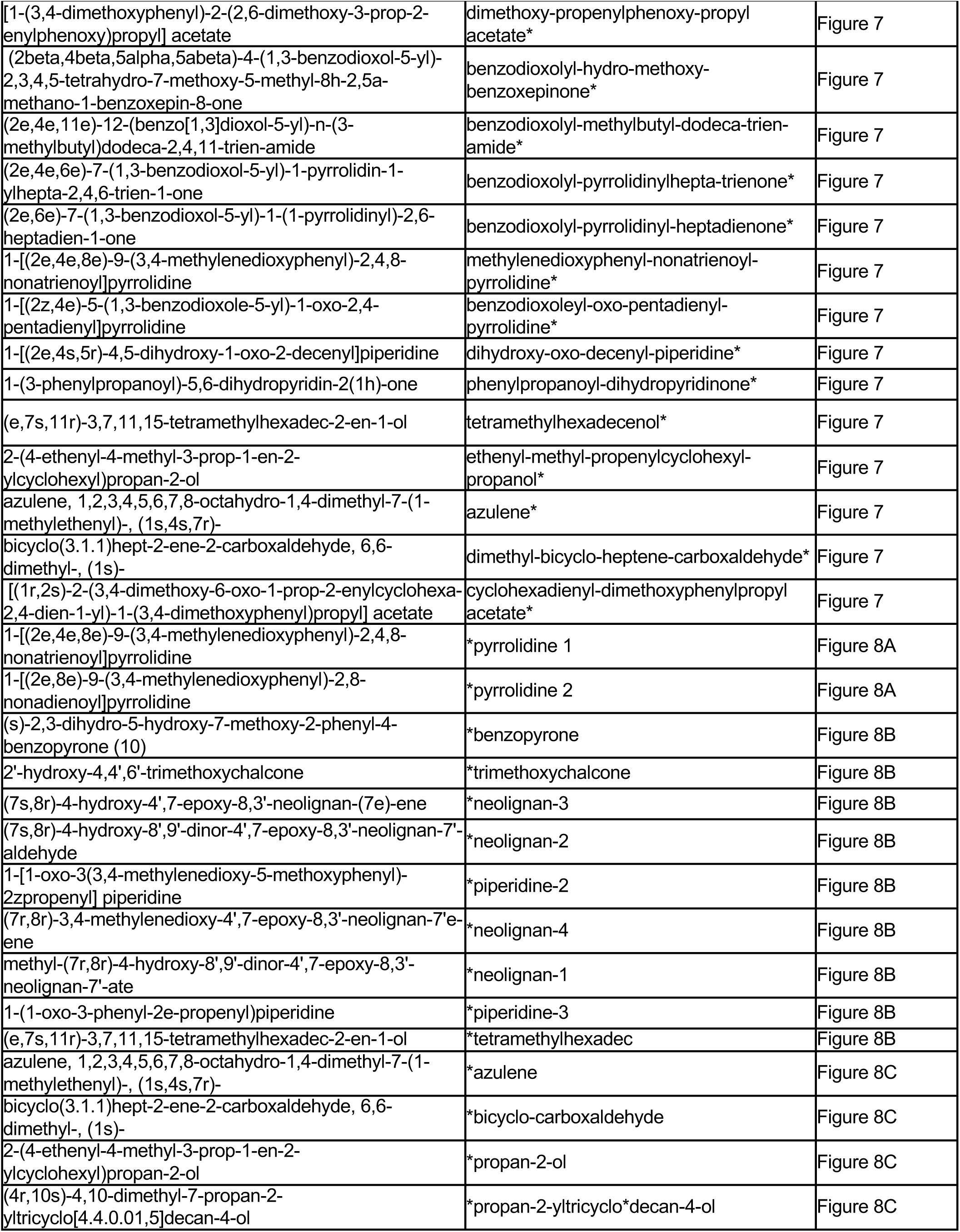
List of full terms for truncated compound names in Figure 4A.

**Supplemental Figure C.**
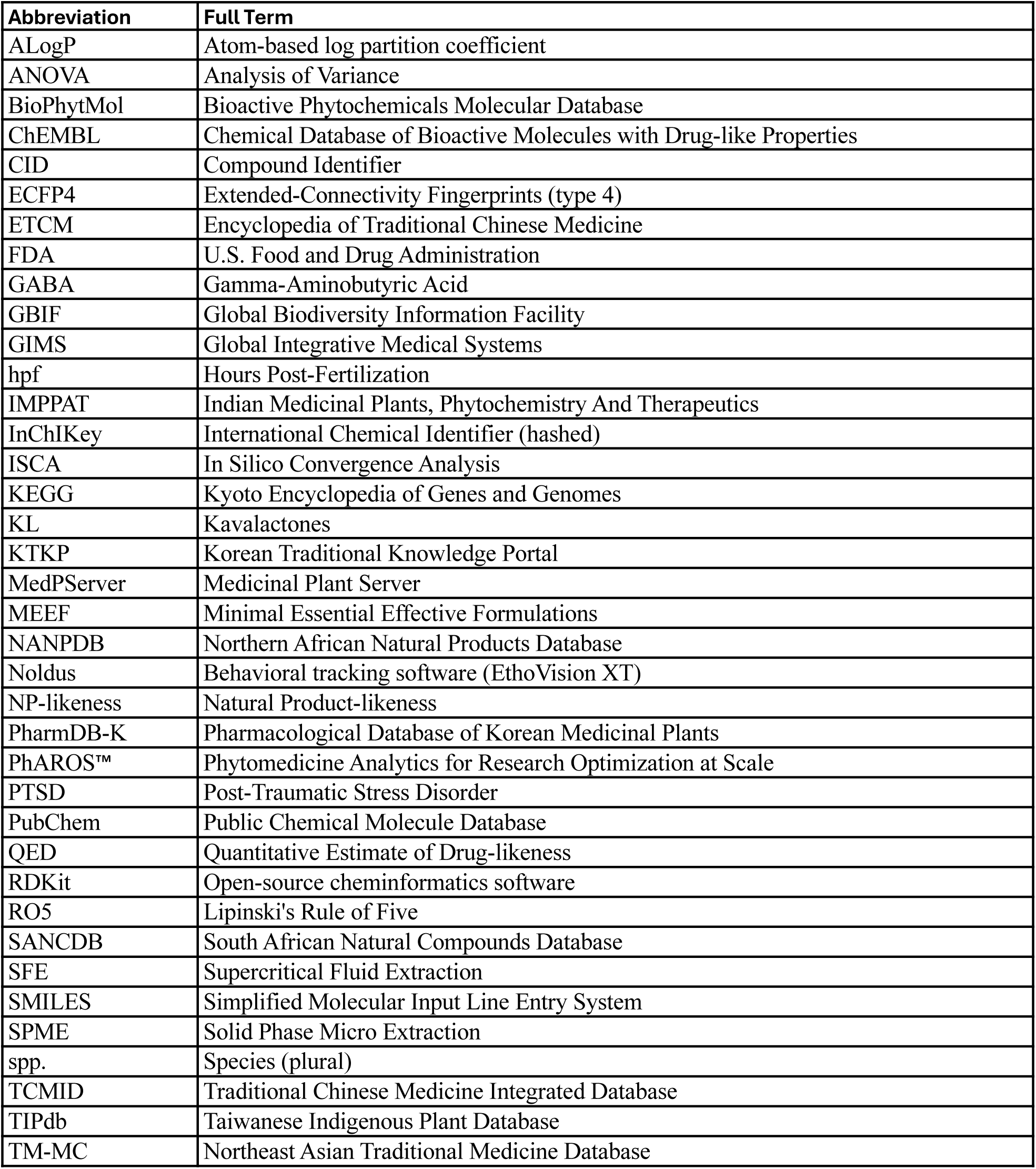
Abbreviation list.

